# ASPP2/PP1 complexes maintain the integrity of pseudostratified epithelia undergoing remodelling during morphogenesis

**DOI:** 10.1101/2020.11.03.366906

**Authors:** Christophe Royer, Elizabeth Sandham, Elizabeth Slee, Jonathan Godwin, Nisha Veits, Holly Hathrell, Felix Zhou, Karolis Leonavicius, Jemma Garratt, Tanaya Narendra, Anna Vincent, Celine Jones, Tim Child, Kevin Coward, Chris Graham, Xin Lu, Shankar Srinivas

## Abstract

During development, pseudostratified epithelia undergo large scale morphogenetic events associated with increased mechanical stress. The molecular mechanisms that maintain tissue integrity in this context are poorly understood. Using a variety of genetic and imaging approaches, we uncover that the ASPP2/PP1 complex ensures proper epiblast and proamniotic cavity architecture via a mechanism that specifically prevents the most apical daughter cells from delaminating apically following cell division events. The ASPP2/PP1 complex achieves this by maintaining the integrity and organisation of the F-actin cytoskeleton at the apical surface of dividing cells. ASPP2/PP1 is also essential during gastrulation in the primitive streak, in somites and in the head fold region, suggesting that this complex is required across a wide range of pseudostratified epithelia during morphogenetic events that are accompanied by intense tissue remodelling and high cell proliferation. Finally, our study also suggests that the interaction between ASPP2 and PP1 is essential to the tumour suppressor function of ASPP2 which may be particularly relevant in the context of tissues that are subject to increased mechanical stress.

## INTRODUCTION

Pseudostratified epithelia are common building blocks and organ precursors throughout embryonic development in a wide array of organisms^1^. As in other epithelia, their cells establish and maintain apical-basal polarity. However, their high nuclear density, high proliferation rate and nuclei movement during interkinetic nuclear migration (IKNM) make them unique. As IKNM proceeds, mitotic cells round up at the apical surface of the epithelium before dividing. During this process, mitotic cells generate enough force to locally distort the shape of the epithelium^2^ or accelerate invagination^3^. During development, pseudostratified epithelia are also subject to large scale morphogenetic events that dramatically affect their shape and organisation. This is particularly true during gastrulation when cells apically constrict in the primitive streak as they push their cell body basally to eventually delaminate into the underlying mesoderm cell layer^4–7^ or when the ectoderm is reshaped to form the head folds. The combined mechanical strains due to IKNM and morphogenetic events poses an incredible challenge for pseudostratified epithelia to maintain tissue integrity during development. However, the molecular mechanisms that allow them to cope with increased mechanical stress are poorly defined.

During these morphogenetic events, epithelial cells continually rely on apical constrictions involving specific F-actin cytoskeleton organisation and actomyosin contractility to modify tissue shape and organisation^8,9^. As cells apically constrict, the coupling of apical junctions to the actomyosin network is essential in transmitting forces across tissues^10^. Reciprocally, as apical constrictions reshape the apical domain of epithelial cells, apical junctions must be able to withstand the forces generated to maintain tissue integrity. Apical-basal polarity components, such as Par3, are vital for apical constrictions^11^ and the integrity of apical junctions^12^. However, it remains unknown how components of the apical-basal polarity machinery maintain tissue integrity in conditions of increased mechanical stress as morphogenetic events occur.

ASPP2 is a Par3 interactor and component of the apical junctions^13,14^. Here, using a variety of genetic and imaging approaches, we show that during morphogenetic events crucial for the normal development of the early post-implantation embryo, ASPP2 maintains epithelial integrity in pseudostratified epithelia under increased mechanical stress. ASPP2 is required for proamniotic cavity formation, the maintenance of primitive streak architecture, somite structure and head fold formation. In the proamniotic cavity, ASPP2 maintains epithelial architecture by preventing apical daughter cells from escaping the epiblast. Mechanistically, we show that this is achieved via the ability of ASPP2 to directly recruit protein phosphatase 1 and through its essential role in maintaining F-actin cytoskeleton organisation at the apical junctions. Our results show that ASPP2 is an essential component of a system that maintains tissue integrity under conditions of increased mechanical stress in a broad range of tissues.

## RESULTS

### The phosphatase and polarity function of ASPP2 are not required for trophectoderm development

ASPP2 can regulate both apical-basal cell polarity and the phosphorylation status of YAP/TAZ through its interaction with Par3^13,14^ and PP1^15,16^ respectively. Both cell polarity and the phosphorylation of YAP and TAZ are crucial to trophectoderm (TE) development^17–21^. We therefore started with the hypothesis that ASPP2 may be important for outside cell polarisation and TE fate determination during preimplantation development. In support of this, we found that ASPP2 could start to be detected as early as E2.5 at cell-cell junctions (Figure S1A). As seen in other examples of polarised epithelia^13,14,16,22^, ASPP2 was strongly localised to the apical junction in the TE from the 32-cell stage onwards (Figure 1A). This localisation pattern was similar in human blastocysts, suggesting that ASPP2 behaves in a similar way across mammals (Figure S1B).

**Figure 1:**
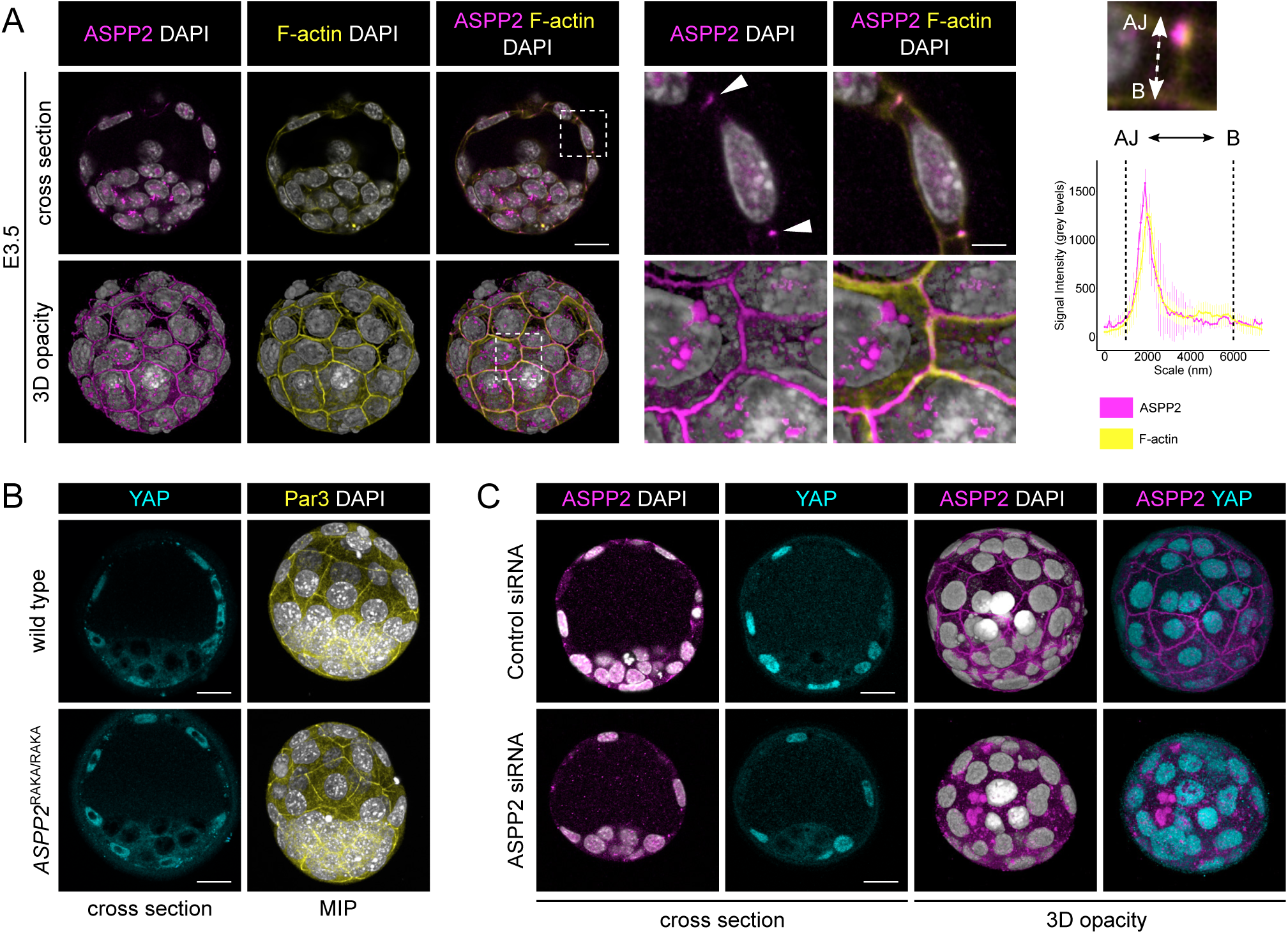
The ASPP2/PP1 complex is not required during preimplantation development. A. ASPP2 was detected by indirect immunofluorescence in E3.5 embryos at the early blastocyst stage to analyse its localisation pattern. A cross section through the equatorial plane of a representative embryo is shown (top row), as well as a 3D opacity rendering of the same embryo (bottom row). The F-actin cytoskeleton and nuclei were visualised using Phalloidin and DAPI respectively. A magnified image of the dashed area is shown on the right. Note how ASPP2 colocalises with F-actin at the apical junctions in cells of the trophectoderm (TE) (white arrowheads). This is quantified in the juxtaposed graph showing ASPP2 and F-actin signal intensity along the apical-basal axis of cell-cell junctions. AJ: apical junction; B: base of the trophectoderm. Scale bars: 20 μm and 5 μm (for the magnification). B. The localisation pattern of YAP and Par3 were analysed in wild type and *ASPP2*^RAKA/RAKA^ embryos by indirect immunofluorescence. A cross section of representative embryos through the equatorial plane shows the localisation of YAP in the nuclei of the TE in both wild type and *ASPP2*^RAKA/RAKA^ embryos. Maximum intensity projections of these embryos show the localisation of Par3 at the level of apical junctions in the TE. Scale bar: 20 μm. C. ASPP2 knockdown in E3.5 embryos using siRNA against ASPP2 mRNA. ASPP2 knockdown was confirmed by indirect immunofluorescence. Note how signal at the apical junctions is specific to ASPP2. Note how YAP is normally localised to the nuclei of TE cells in ASPP2-depleted embryos. Scale bar: 20 μm.

A Previous study revealed that ASPP2 may play a role during early embryogenesis, as ASPP2-mutant embryos in which exons 10-17 were deleted could not be recovered at E6.5^23^. However, the phenotype of these embryos was not described, and earlier stages were not examined. To investigate the role of ASPP2 during preimplantation development we generated *ASPP2*-null embryos in which exon 4 was deleted (*ASPP2*^ΔE4/ΔE4^) resulting in a frameshift and early stop codons. To be able to distinguish between phenotypes relating to ASPP2’s PP1 regulatory function and other functions, we also generated embryos homozygous for a mutant form of ASPP2 that was specifically unable to associate with PP1 (*ASPP2*^RAKA/RAKA^)^24^. *ASPP2*^ΔE4/ ΔE4^ and *ASPP2*^RAKA/RAKA^ blastocysts appeared to be morphologically normal with properly formed blastocyst cavities (Figure 1B and S1C). YAP was clearly nuclear in the TE and cytoplasmic in the inner cell mass (ICM) suggesting that the ICM and TE lineage were properly allocated (Figure 1B). The polarity protein Par3 (Figure 1B) and F-actin cables (Figure S1C) were strongly localised at the apical junctions, suggesting that polarity and overall cell architecture were normal. It was sometimes possible to see some residual ASPP2 protein at the apical junction in early *ASPP2*^ΔE4/ΔE4^ blastocysts, potentially due to residual maternal ASPP2 expression. To eliminate the possibility that perdurance of maternally encoded ASPP2 compensates for the zygotic mutations, we microinjected 1-cell embryo with siRNA against ASPP2 and cultured them to the blastocyst stage. Control and ASPP2-depleted embryos were morphologically indistinguishable. The localisation of YAP was similar between control and ASPP2-depleted embryos (Figure 1C). YAP phosphorylation at Serine 127 was stronger in the cytoplasm of ICM cells in comparison to the TE in both controls and ASPP2-depleted embryos (Figure S1D). Taken together, these results show that neither ASPP2’s polarity function nor its PP1 regulatory function are required during preimplantation development.

### ASPP2/PP1 interaction is required for proamniotic cavity architecture

Since it has previously been shown that deletion of ASPP2 may be embryonic lethal around E6.5^23^, we next investigated whether ASPP2 was required at early post-implantation stages. To test this, we generated *ASPP2*^ΔE4/ΔE4^ embryos at different stages and examined the localisation of Par6 and F-actin to assess apical-basal polarity and overall tissue organisation, respectively. At E5.5, *ASPP2*^ΔE4/ΔE4^ embryos did not exhibit obvious morphological defects in comparison to wild type and heterozygous litter mates. Polarised Par6 could be detected at the apical membrane in the visceral endoderm (VE) and the epiblast, suggesting that both cell layers properly polarised (Figure S2A). In contrast, E6.5 *ASPP2*^ΔE4/ΔE4^ embryos exhibited strong morphological defects in comparison to wild type and heterozygous litter mates. The proamniotic cavity either entirely failed to form (6 embryos out of 10 mutants) or was greatly reduced in size at E6.5 (4 embryos out of 10 mutants) (Figure 2A) and always absent at E7.5 (Figure S2B). In embryos lacking a proamniotic cavity, the epiblast was disorganised, and instead of being a pseudostratified epithelium, appeared multi-layered. This seemed to be the result of an ectopic accumulation of cells from the epiblast in place of the proamniotic cavity. The ectopic cells in the centre of the embryo exhibited a complete lack of polarised Par6 and in embryos with reduced cavity size, the accumulated cells showed reduced apical Par6 (Figure 2B). This suggested that the ectopic accumulation of cells where the proamniotic cavity ought to be was accompanied by a progressive loss of cell polarity in the epiblast. F-actin localisation was also profoundly abnormal in these cells (Figure 2B). In wild type embryos, F-actin was enriched at apical junctions, whereas in *ASPP2*^ΔE4/ΔE4^ embryos it was distributed more uniformly across the apical surface (Figure 2C). This suggests that ASPP2 is required for organising the F-actin cytoskeleton at the apical junctions.

**Figure 2:**
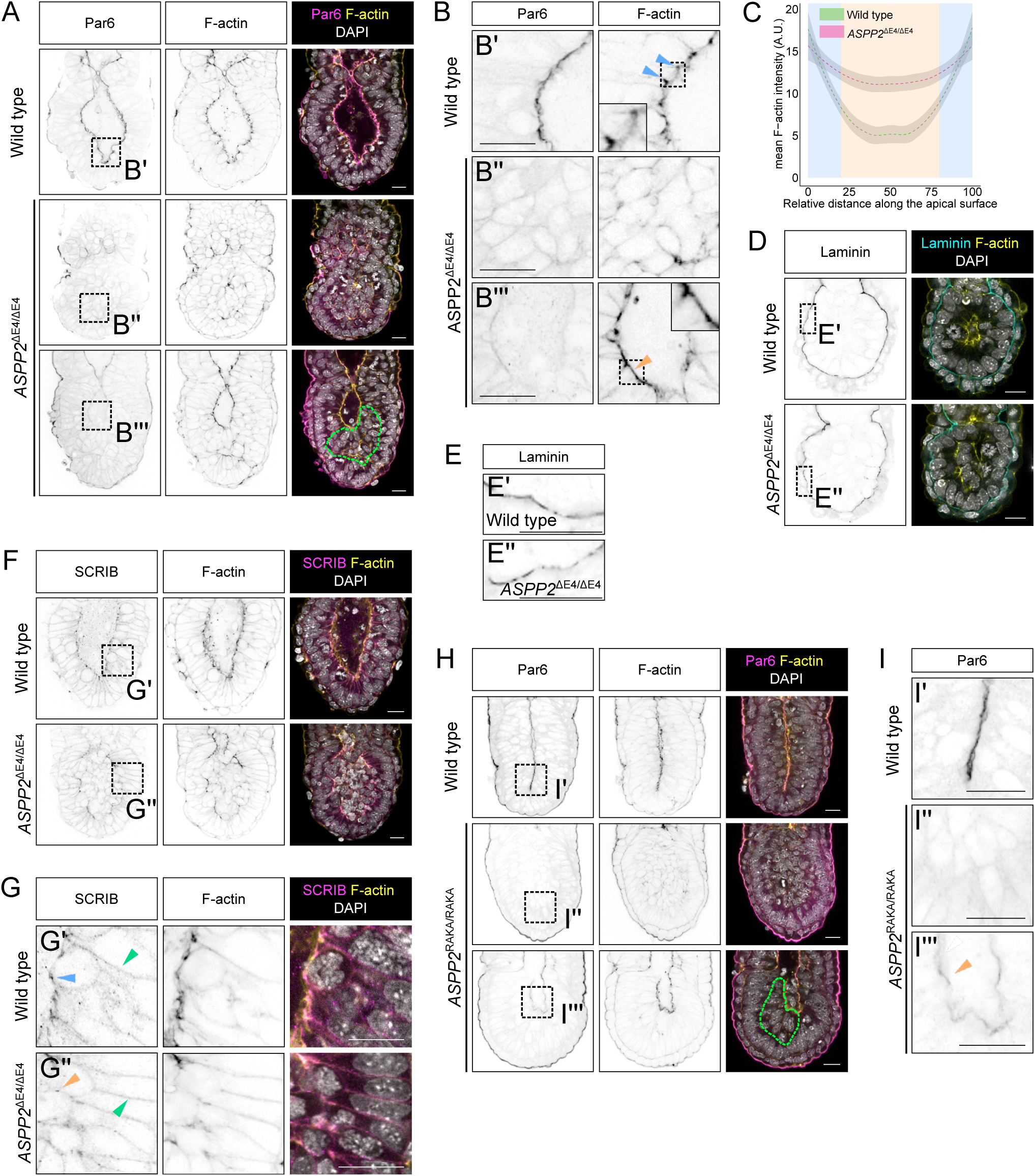
The ASPP2/PP1 complex is required for the formation of the proamniotic cavity. A. Immunofluorescence of wild type and *ASPP2*^ΔE4/ΔE4^ E6.5 embryos using an anti-Par6 antibody. The phenotypic variability of *ASPP2*^ΔE4/ΔE4^ embryos is illustrated, with embryos either lacking cavities (middle row) or exhibiting smaller cavities (bottom row). The green dashed line highlights the ectopic accumulation of cells in the epiblast of *ASPP2*^ΔE4/ΔE4^ embryos. B. Magnification of the corresponding regions shown in panel A. Blue arrowheads highlight the enrichment of F-actin at apical junctions in the epiblast. Note how F-actin is not enriched at apical junctions but is instead more homogenously distributed across the apical surface of epiblast cells in *ASPP2*^ΔE4/ΔE4^ embryos (orange arrowhead). The insets within images are 2x magnifications of the corresponding dashed areas. C. Quantification of F-actin signal intensity along the apical surface of epiblast cells of wild type (n=3 embryos, 5 measurements per embryo) and *ASPP2*^ΔE4/ΔE4^ embryos (n=3 embryos, 5 measurements per embryo). Measurements were made on cross sections along the apical domain of individual epiblast cells from apical junction to apical junction (represented with a blue background in the graph). See material and methods for details. D. Immunofluorescence of wild type and *ASPP2*^ΔE4/ΔE4^ E5.5 embryos using an anti-Laminin antibody. E. Magnification of the corresponding dashed areas in panel D. F. Immunofluorescence of wild type and *ASPP2*^ΔE4/ΔE4^ E6.5 embryos using an anti-SCRIB antibody. G. Magnification of the corresponding dashed areas in panel F. Green arrowheads highlight basolateral SCRIB. Note the enrichment of SCRIB at the apical junctions in the epiblast of wild type embryos (blue arrowhead) and its absence in the corresponding localisation in *ASPP2*^ΔE4/ΔE4^ embryos (orange arrowhead). H. Immunofluorescence of wild type and *ASPP2*^RAKA/RAKA^ E6.5 embryos using an anti-Par6 antibody. The green dashed line highlights the ectopic accumulation of cells in the epiblast of *ASPP2*^RAKA/RAKA^ embryos. I. Magnification of the corresponding dashed regions in H. Note the reduced amount of Par6 along the apical domain of epiblast cells in *ASPP2*^RAKA/RAKA^ embryos (orange arrowhead). Nuclei and the F-actin cytoskeleton were visualised with DAPI and Phalloidin respectively. Scale bars: 20 μm.

Because signals from the basement membrane are believed to be essential for proamniotic cavity formation^25^, we examined the localisation of laminin and found it unaltered in *ASPP2*^ΔE4/ΔE4^ embryos. This suggested that the loss of cell polarity and ectopic accumulation of cells was not accompanied by, or due to, breakage in the basement membrane (Figure 2D and 2E). Finally, when we examined the basolateral membrane marker SCRIB, we observed that its basolateral localisation was unaffected in *ASPP2*^ΔE4/ ΔE4^ embryos. However, SCRIB was also strongly expressed at the apical junctions in the epiblast of wild type embryos. This particular localisation pattern was intermittently disrupted in *ASPP2*^ΔE4/ΔE4^ embryos, specifically at the interface between cells of the epiblast and cells ectopically accumulating in the proamniotic cavity (Figure 2F and 2G). Together, these results suggest that the apparent loss of apical cell polarity seen is *ASPP2*^ΔE4/ΔE4^ embryos originates from defects specific to the apical junctions rather than at the level of the basolateral or basement membranes.

The outside VE monolayer epithelium also normally expresses ASPP2 (Figure 5A and S5A and B) but was intact and exhibited apical Par6 in *ASPP2*^ΔE4/ ΔE4^ mutants (Figure 2A), suggesting that its epithelial architecture and integrity was maintained and that ASPP2 is required specifically in the epiblast at this stage. To verify that an epiblast specific requirement for ASPP2 led to the failure of proamniotic cavity formation in *ASPP2*^ΔE4/ΔE4^ embryos, we conditionally ablated ASPP2 expression in just the epiblast (*ASPP2*^EpiΔE4/ΔE4^ embryos) (Figure S2C). These embryos phenocopied *ASPP2*^ΔE4/ΔE4^ embryos, demonstrating that at this stage, ASPP2 is required only in the epiblast.

To test whether this requirement for ASPP2 is rooted in its ability to recruit and regulate PP1, we analysed *ASPP2*^RAKA/RAKA^ embryos at E6.5. We found that *ASPP2*^RAKA/RAKA^ embryos, similarly to *ASPP2*^ΔE4/ΔE4^ embryos, exhibit either reduced proamniotic cavity size or no cavity at all. This was again accompanied by a reduced apical Par6 in the epiblast when the proamniotic cavity was of reduced size and absence of apical Par6 when no cavity was present (Figure 2H and 2I). The basolateral localisation of SCRIB once again was not affected, whereas its localisation at apical junctions was severely disrupted in *ASPP2*^RAKA/RAKA^ embryos (Figure S2D and S2E). This shows that at E6.5, *ASPP2*^RAKA/ RAKA^ mutant embryos have an identical phenotype to *ASPP2*^ΔE4/ΔE4^ mutant embryos, demonstrating the key role of the ASPP2/PP1 interaction in regulating epiblast and proamniotic cavity architecture.

### ASPP2 controls apical daughter cell reincorporation into the epiblast

Our results so far show that ASPP2 is essential for the architecture of the epiblast and the formation of the proamniotic cavity. When ASPP2’s function is impaired, apolar cells accumulate ectopically in place of the proamniotic cavity. However, it remained unclear how this occurred and what biological process ASPP2 actually controls in the epiblast. Amongst possible explanations is that the phenotype was the consequence of epiblast cells delaminating apically into the proamniotic cavity because of a drastic shift in the proportion of orthogonal cells divisions, a breakdown of the apical junction domain, a failure of daughter cells reincorporating basally following cell divisions or a combination of these. To answer this question, we generated *ASPP2*^ΔE4/ΔE4^ embryos with fluorescently labelled membranes, which enabled us to follow the movement of epiblast cells in these embryos by time-lapse confocal microscopy (Figure 3A and Movie 1). In wild type and heterozygous embryos, we could observe the movement of cell bodies along the apical-basal axis during interkinetic nuclear migration (IKNM), with mitotic cells rounding up at the apical surface of the epiblast before dividing as previously described^26^.

**Figure 3:**
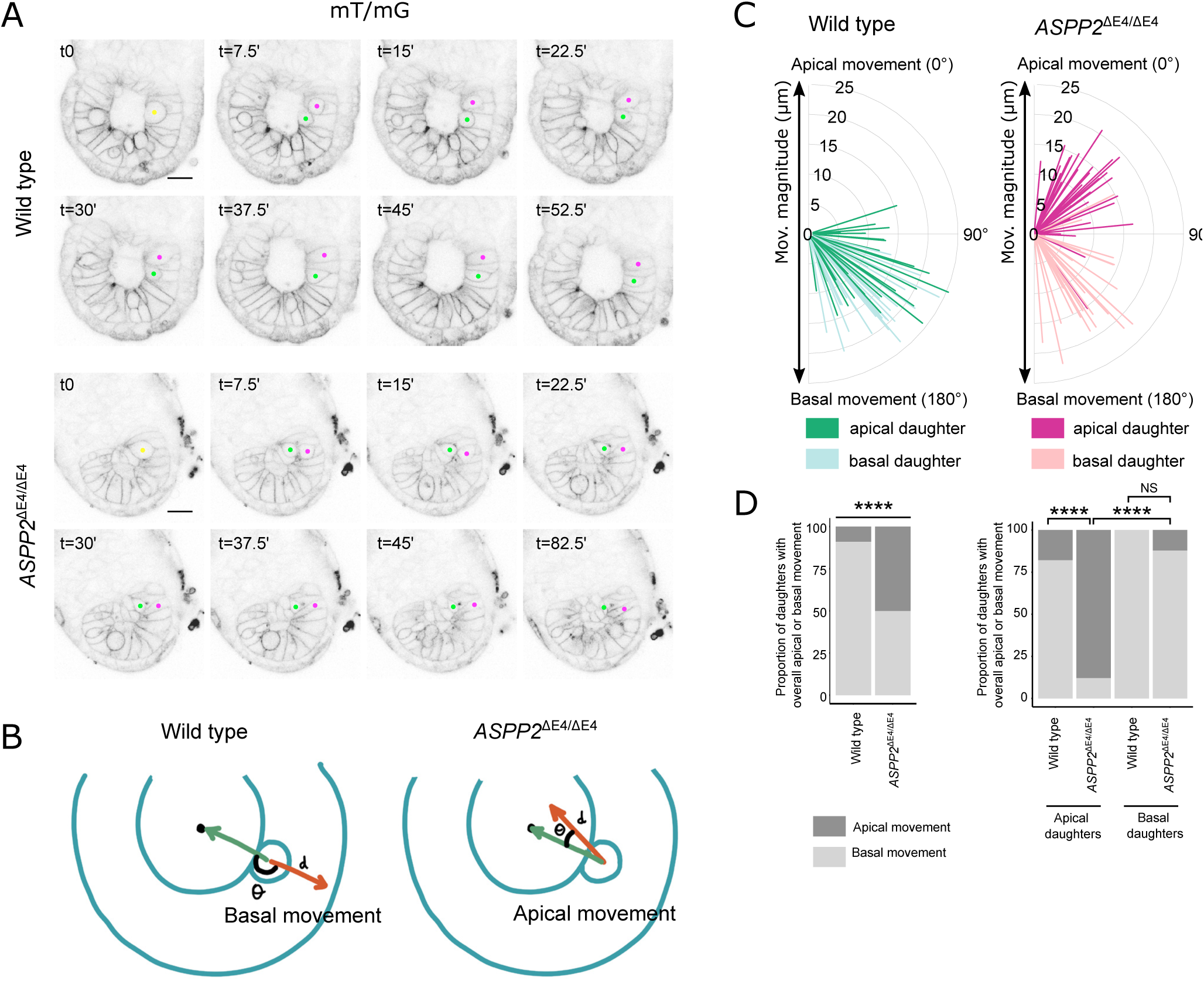
ASPP2 is required for apical daughter cell reincorporation into the epiblast following cell division events. A. Time lapse imaging of wild type and *ASPP2*^ΔE4/ΔE4^ embryos. mT/mG-labelled cell membranes were used to manually track cell movement. Yellow dots highlight mother cells at the apical surface of the epiblast immediately prior to a cell division event. Green and magenta dots identify the resulting daughter cells. Note how both daughters reintegrate the epiblast in the wild type whereas one of the two daughters fails to do so in the absence of ASPP2 even after a prolonged period of time (t=82.5’). B.Diagram illustrating the method used to quantify daughter cell movement following cell divisions. Daughter cell movement was characterised by both the distance travelled (d) and the direction of travel (θ) expressed as the angle between the reference vector (the green vector starting from the initial position of the mother cell prior to the division event to the centre of the embryonic region) and the vector characterising absolute daughter cell movement (the red vector starting from the initial position of the mother cell prior to the division event to the final position of the daughter cell). The left panel illustrates the case of a daughter moving basally to reincorporate the epiblast and the right panel describes abnormal daughter cell movement towards the centre of the embryonic region such as seen in *ASPP2*^ΔE4/ΔE4^ embryos. C. Graph quantifying cell movement in wild type (n=3 embryos, 56 cells) and *ASPP2*^ΔE4/ΔE4^ embryos (n=3 embryos, 66 cells). For a given pair of daughter cells, each daughter was defined as “apical” or “basal” depending on their respective position relative to the centre of the embryonic region immediately after a cell division event. D. Proportion of daughter cells with an overall apical or basal movement in wild type and *ASPP2*^ΔE4/ΔE4^ embryos. Left panel: Quantification of the proportion of daughter cells with an overall apical (θ from 0° to 90°) or basal movement (θ from 90° to 180°) in wild type and *ASPP2*^ΔE4/ΔE4^ embryos. Right panel: quantification of the proportion of apical and basal daughters with an overall apical (θ from 0° to 90°) or basal movement (θ from 90° to 180°) in wild type and *ASPP2*^ΔE4/ΔE4^ embryos. **** p<0.0001, NS: non-significant (Fisher’s exact test of independence).

We first analysed the orientation of cell divisions but could not detect differences in overall cell division angle in *ASPP2*^ΔE4/ΔE4^ embryos in comparison to controls (Figure S3A), even when division events were binned into categories as “orthogonal”, “parallel” or “oblique”. To determine if there was a defect in IKNM, we also analysed the distance at which cell divisions occurred from the basement membrane of the epiblast. Again, there was no notable difference between *ASPP2*^ΔE4/ΔE4^ and wild type embryos, suggesting that even in the absence of apical-basal polarity and the proamniotic cavity, cells of the epiblast were able to proceed with IKNM (Figure S3B).

We next looked at the behaviour of daughter cells after cytokinesis. In wild type embryos, following cell divisions, the cell body of both daughters moved basally so that they came to span the entire height of the epithelium along the apical-basal extent of the epiblast. In contrast, in *ASPP2*^ΔE4/ΔE4^embryos, dividing cells moved towards the embryonic centre as normal, but upon division, daughter cells delaminated apically towards the centre of mass of the embryonic region (Figure 3A). This suggested that ASPP2 may be specifically required for the retention of daughter cells within the epiblast. To further characterise this failure of dividing cells to reintegrate into the epiblast epithelium, we quantified the movement of daughter cells from the initial point of cell division (Figure 3B-D). We found that in wild type and heterozygous embryos, this movement was almost always basal for both daughters (51/56, 91.1%). In contrast, in mutant embryos, for half the daughter cells (33/66, 50%) the movement was apical (Figure 3C and 3D). We found that in the majority of cases (29/33, 87.9%), it was the daughter that was relatively more apically positioned with respect to its sister that abnormally moved apically following cell divisions (Figure 3D). This suggests that ASPP2 is involved in a novel mechanism specifically required for apical daughter cell reintegration into the pseudostratified epiblast following cell division, which is crucial in maintaining the architecture of the epiblast and proamniotic cavity.

### ASPP2/PP1 complexes maintain epithelial integrity in regions of high mechanical stress

Our results all point to an important role for ASPP2 in regulating tissue architecture, possibly via the regulation of F-actin organisation at the apical junction. However, this was difficult to study in *ASPP2*^ΔE4/ΔE4^ and *ASPP2*^RAKA/RAKA^ embryos on a C57BL/6 background because of the relative severity of the defect. We therefore bred *ASPP2*^RAKA/RAKA^ mutation into a BALB/c background, to take advantage of the fact that ASPP2 phenotypes are often not as dramatic in this background^14^. Consistent with this, BALB/c *ASPP2*^RAKA/ RAKA^ homozygous embryos completely bypassed the phenotype at E6.5 observed in C57BL/6 *ASPP2*^RAKA/ RAKA^ embryos. Instead, the phenotype of these embryos was milder, and they were only grossly different from wild type and heterozygous embryos one day later, at E7.5. *ASPP2*^RAKA/RAKA^ embryos exhibited two distinct phenotypes. The majority (34/41, 82.9%), that we termed type I embryos, exhibited a strong accumulation of cells in their posterior, suggestive of a defect in the primitive streak (Figure S4A and 4A). A minority (7/41, 17.1%), that we termed type II embryos, were developmentally delayed but did not exhibit any structural defects.

Given that *ASPP2*^ΔE4/ΔE4^ and *ASPP2*^RAKA/RAKA^ in a Bl/6 background showed striking abnormalities in the localisation of F-actin, we examined the localisation of F-actin in the posterior of type I *ASPP2*^RAKA/RAKA^ embryos at E7.5 (Figure 4B and Figure S4C). In wild type embryos, we were able to clearly identify cells apically constricting and pushing their cell body basally towards the nascent mesodermal cell layer. This is characteristic of cells in the primitive streak in the process of delaminating basally^4^. It was also evident that F-actin was enriched at the apical junctions in these cells (Figure 4C). In contrast, *ASPP2*^RAKA/RAKA^ embryos exhibited a clear ectopic accumulation of cells apical to the primitive streak as visualised by T expression (Figure 4B). In these cells, F-actin was abnormally uniformly disturbed along the apical surface, with no clear apical-junction enrichment (Figure 4C). To investigate this further, we performed Airyscan super-resolution imaging of these embryos (Figure 4D-F).This revealed that the mesh-like structure normally formed by F-actin at the apical junctions in cells of the epiblast was severely disrupted in the posterior of *ASPP2*^RAKA/ RAKA^ embryos. Instead, F-actin appeared to form spike-like structures at the surface of cells in this region (Movie 2) (figure 4E and 4F). This profound disruption of F-actin localisation indicates that the PP1 regulatory function of ASPP2 is required for F-actin organisation in the cells of the primitive streak.

**Figure 4:**
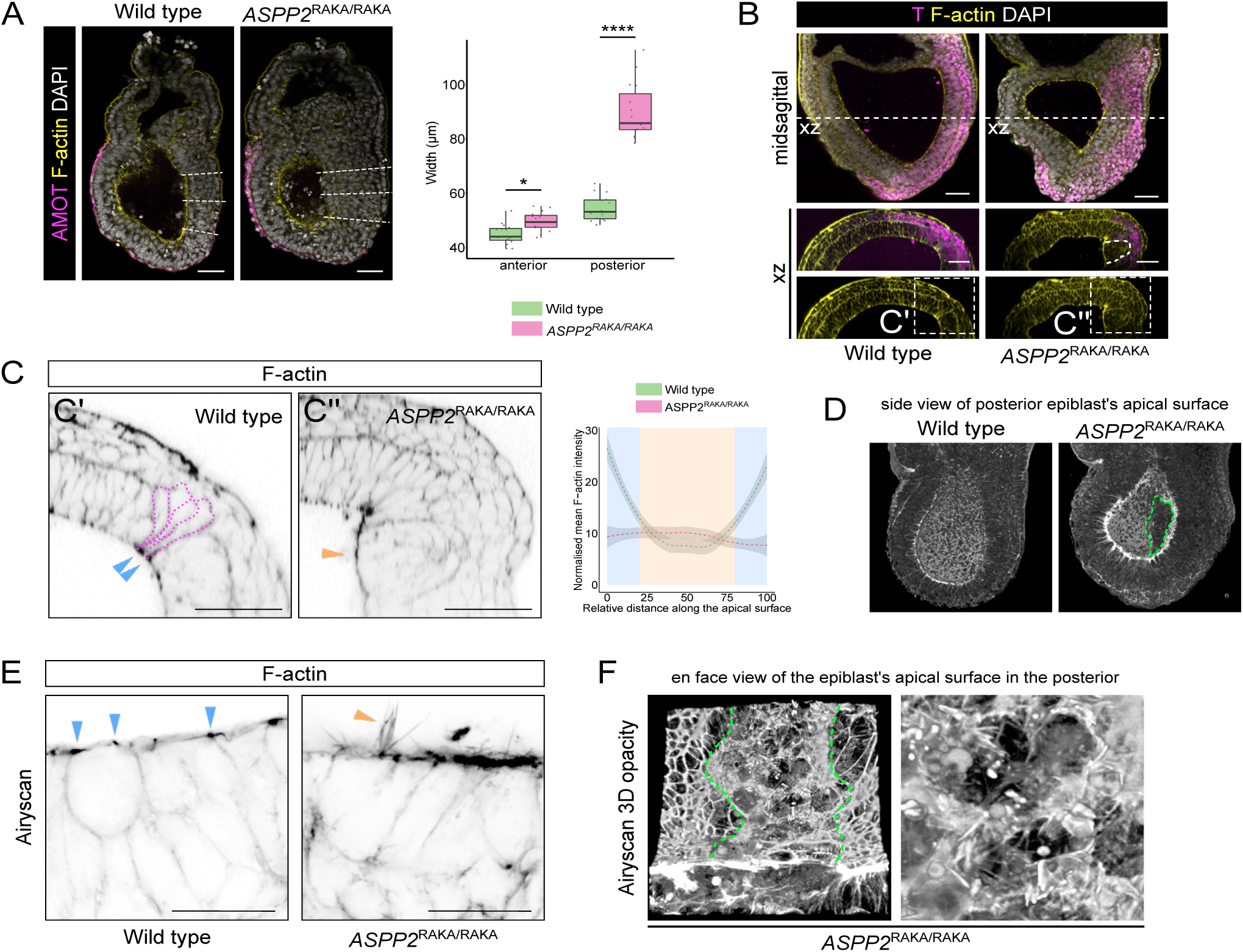
ASPP2 is required for epithelial integrity in the primitive streak. A. Posterior thickening in E7.5 *ASPP2*^RAKA/RAKA^ embryos in a BALB/C background. Left panel: the anteroposterior axis was defined using AMOT localisation pattern. Right panel: comparison of tissue thickness in the anterior (3 measurements per embryo) and the posterior (3 measurements per embryo) of wild type (n=5 embryos) and *ASPP2*^RAKA/RAKA^ embryos (n=5 embryos). * p<0.05, ****p<0.0001 (nested ANOVA). B. Cells accumulate in the primitive streak region of *ASPP2*^RAKA/RAKA^ embryos. Immunofluorescence of E7.5 wild type and *ASPP2*^RAKA/RAKA^ embryos using a T (Brachyury) antibody. C. Cells ectopically accumulating in the primitive streak region are unable to apically constrict and do not have enriched F-actin at the apical junctions (orange arrowhead) in comparison to wild type (blue arrow heads). Magenta dotted lines highlight cells apically constricting to push their cell bodies basally towards the underlying mesoderm cell layer. Right panel: quantification of F-actin signal intensity along the apical surface of epiblast cells in the primitive streak region of wild type (n=3 embryos, 5 cells per embryo) and *ASPP2*^RAKA/RAKA^ embryos (n=3 embryos, 5 cells per embryo). Measurements were made on cross sections along the apical domain of individual epiblast cells from apical junction to apical junction (represented with a blue background in the graph). D-F. Airyscan imaging reveals the extent of F-actin disorganisation at the surface of cells accumulating ectopically in the primitive streak region of *ASPP2*^RAKA/ RAKA^embryos. D. 3D opacity rendering of embryo optical halves, enabling visualisation of the apical surface of epiblast cells in the proamniotic cavity. Note the absence of the typical F-actin mesh pattern at the apical surface of cells in the posterior of *ASPP2*^RAKA/RAKA^ embryos (green dotted line). E. F-actin localisation pattern in the primitive streak region of wild type and *ASPP2*^RAKA/RAKA^ embryos. F-actin was enriched at the apical junctions of wild type embryos (blue arrowheads). Orange arrowheads highlight the formation of F-actin spike-like structures at the contact-free surface of *ASPP2*^RAKA/RAKA^ embryos. F. En face view of the epiblast’s apical surface in the posterior of an *ASPP2*^RAKA/RAKA^ embryo. Green dotted lines demarcate the disorganised apical region of the posterior and the more organised lateral regions of the epiblast. Right panel: magnification of the epiblast’s apical surface in the posterior of an *ASPP2*^RAKA/RAKA^ embryo showing F-actin forming spike-like structures. Nuclei and the F-actin cytoskeleton were visualised with DAPI and Phalloidin respectively. Scale bars: 20 μm.

The epiblast specific requirement for ASPP2 despite its broad expression (Figure 1A and 6A-C) and the localisation of the phenotype primarily to the posterior region of the epiblast in the BALB/c background suggest that specific epithelia or epithelial regions are more sensitive to ASPP2 deficiency than others. We hypothesised therefore that ASPP2 may be important particularly for dividing cells to reintegrate within epithelia subject to increased mechanical stress at the apical junction, for example during the apical curving required to form a cavity. In support of this hypothesis, phospho-myosin levels were higher in dividing cells in the epiblast, in particular at the apical junctions. This shows that actomyosin contractility increases at the apical junctions of dividing cells, which might result in increased mechanical stress (Figure 5A). One prediction of this hypothesis is that increasing the mechanical stress in Type I *ASPP2*^RAKA/RAKA^ embryos in a BALB/c background might induce an earlier or more severe phenotype reminiscent to that seen in the C57BL/6 background. To test this prediction, we cultured E6.5 BALB/c wild type and mutant embryos (a day before any phenotype is evident in mutants) within the confines of cylindrical cavities made of biocompatible hydrogels, in order to alter their shape^27^ and subject the epiblast epithelium to higher levels of mechanical stress (Figure 5B). Wild type embryos elongated without showing any sign of disrupted tissue integrity (0 out of 4 embryos). Conversely, *ASPP2*^RAKA/RAKA^ embryos showed reduced cavity size with a clear accumulation of cells (2 out of 2 embryos), phenocopying *ASPP2*^ΔE4/ΔE4^ and *ASPP2*^RAKA/ RAKA^ mutant embryos in a C57BL/6 background (Figure 5C). Interestingly, F-actin and Myosin were abnormally distributed at the apical surface of cells accumulating ectopically in *ASPP2*^RAKA/RAKA^ mutant embryos, suggesting that actomyosin contractility was disrupted. Together, This indicates that although *ASPP2*^RAKA/ RAKA^ mutants in a BALB/c background can bypass the proamniotic cavity phenotype, increasing mechanical stress is sufficient to make them again susceptible to it and suggests that ASPP2 may be required in response to increased mechanical stress to maintain epithelial tissue integrity.

### ASPP2 maintains the architecture of the F-actin cytoskeleton during cell division events via its PP1 regulatory function

To understand how the absence of ASPP2 specifically regulates apical daughter cell reintegration into the epiblast, we analysed in detail its localisation pattern. ASPP2 was localised at the apical junctions in the VE (Figure 6A and Figure S5A and B) and the epiblast (Figure 6B and see Fig S2C for antibody specificity). In the former, ASPP2 was uniformly distributed along the apical junctions forming a regular mesh at the surface of the embryo (Figure 6A and Figure S5A and B). In the epiblast however, ASPP2 appeared enriched at specific locations along the apical junctions (Figure 6B). The high curvature of the inner apical surface of the epiblast makes it difficult to examine from standard confocal volumes. We therefore computationally ‘unwrapped’^28^ the apical surfaces of the VE and epiblast so that we could more directly compare them (Figure 6C). This revealed that, although ASPP2 was uniform in its distribution along all junctions in the VE, in the epiblast, it was enriched specifically at F-actin-rich tricellular junctions.

**Figure 5:**
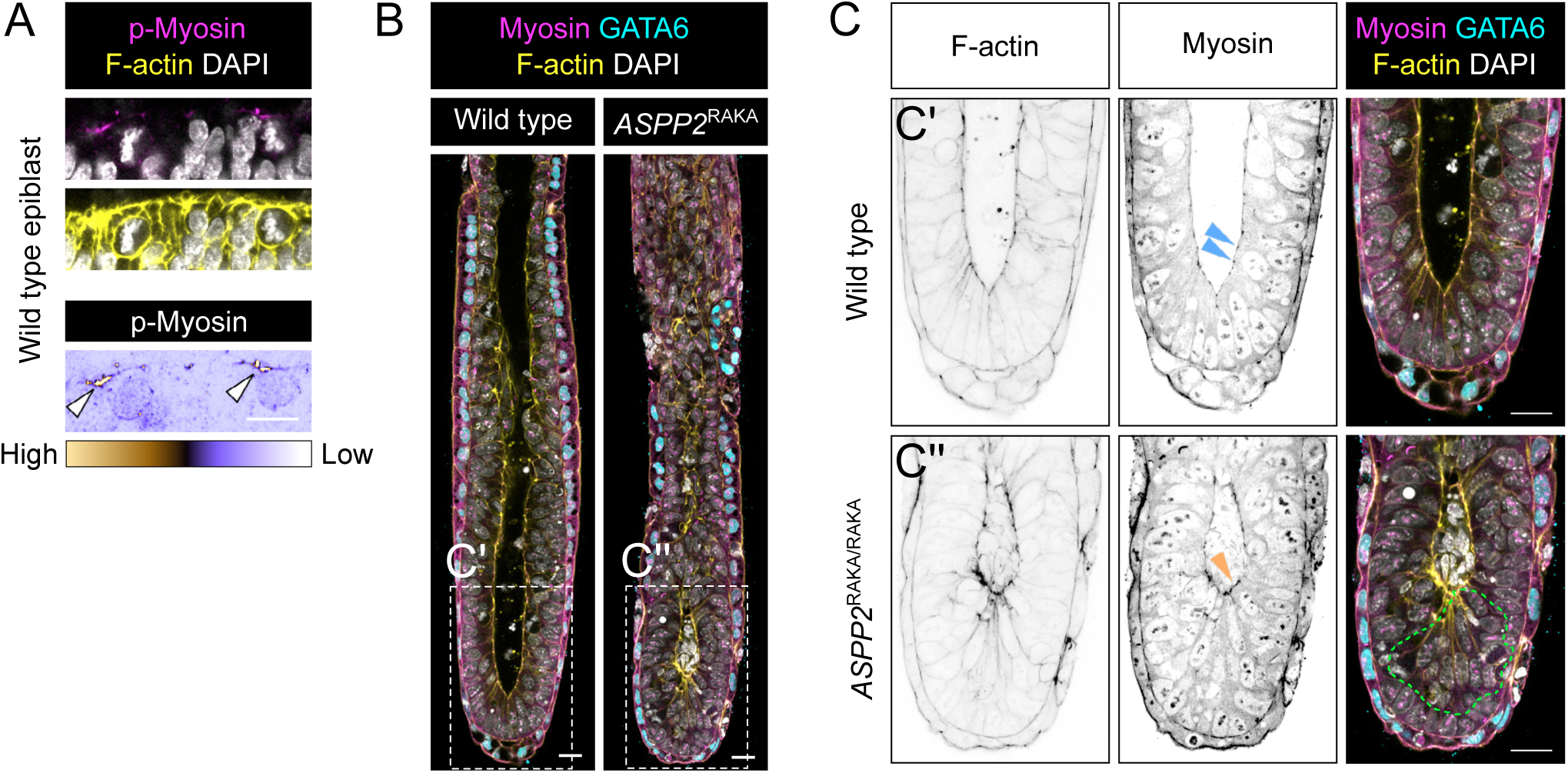
ASPP2^RAKA/RAKA^ embryo are more susceptible to mechanical stress. A. Phospho-myosin levels are higher at the apical junctions of dividing cells in the epiblast (white arrowheads). B. wild type (n=3) and *ASPP2*^RAKA/RAKA^ (n=2) embryos were grown for 30’ in cylindrical cavities made of biocompatible hydrogels. The localisation pattern of GATA6 and Myosin was then analysed by immunofluorescence. C. Magnification of the embryos shown in E. The green dotted line highlights the ectopic accumulation of cells seen in *ASPP2*^RAKA/RAKA^ embryos. The orange arrowhead points to the abnormal distribution of Myosin at the apical surface of these cells. Nuclei and the F-actin cytoskeleton were visualised with DAPI and Phalloidin respectively. Scale bars: 20 μm.

**Figure 6:**
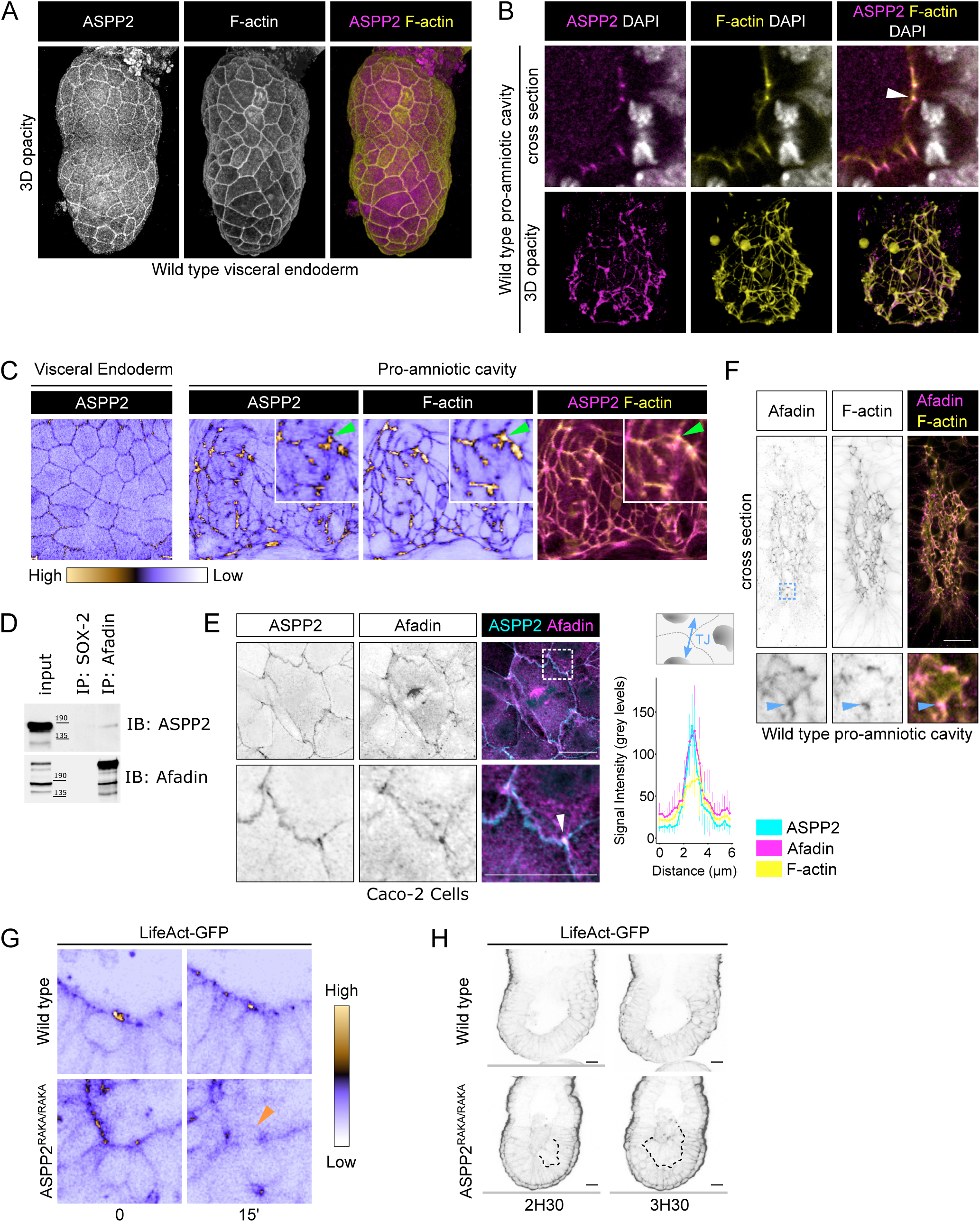
the ASPP2/PP1 complex is required for F-actin organisation during cell division events. A. 3D opacity rendering showing the localisation of ASPP2 in E5.5 wild type embryos at the apical junctions of the visceral endoderm where it colocalises with F-actin. B. Cross section (top row) and 3D opacity rendering (bottom row) of the proamniotic cavity showing the localisation pattern of ASPP2 and F-actin at the apical junctions (white arrowhead). C. The outer surface of the VE and apical surface of the epiblast were computationally ‘unwrapped’, revealing the enrichment of ASPP2 and F-actin at tricellular junctions in the epiblast (green arrowheads). D. The interaction between endogenous ASPP2 and the F-actin-binding protein Afadin was examined in Caco-2 cells by co-immunoprecipitation. E. The localisation pattern of endogenous ASPP2 and Afadin in Caco-2 cells was examined by immunofluorescence. The bottom row represents the magnified region highlighted by a dotted box and shows the enrichment of ASPP2 and Afadin at tricellular junctions. ASPP2, Afadin and F-actin signal intensity was quantified across tricellular junctions (graph on the right). F. The localisation pattern of Afadin in the proamniotic cavity was analysed by immunofluorescence in E6.5 wild type embryos. Afadin colocalised strongly with F-actin at tricellular junctions (blue arrowhead). G. The localisation pattern of F-actin was analysed by time-lapse microscopy in wild type and *ASPP2*^RAKA/RAKA^ LifeAct-GFP positive embryos. Note how apical F-actin is disrupted in *ASPP2*^RAKA/RAKA^ LifeAct-GFP positive embryos following a cell division event (orange arrowhead). H. At later time points, the ectopic accumulation of cells in the epiblast of *ASPP2*^RAKA/RAKA^ LifeAct-GFP positive embryos was evident (dotted line). Nuclei and the F-actin cytoskeleton were visualised with DAPI and Phalloidin respectively. Scale bars: 20 μm.

The enrichment of ASPP2 in regions of high F-actin and the disruption of F-actin localisation in mutants suggests that ASPP2 may somehow be linked to the F-actin cytoskeleton. Because ASPP2 does not possess any known F-actin binding domain, we looked if previously identified ASPP2 binding partners could provide this link. Interestingly, ASPP2 has been found to interact with Afadin in a number of proteomic studies^29,30^. Afadin is an F-actin-binding protein that has previously been shown to not only be enriched at tricellular junctions but to also regulate their architecture^31^. Moreover, at E7.5, *Afadin*-null embryos display a phenotype reminiscent of the phenotype observed in E7.5 *ASPP2*^ΔE4/ΔE4^ embryos (Figure S2B) with cells accumulating in the proamniotic cavity^32^, suggesting that Afadin and ASPP2 have overlapping functions. To confirm that ASPP2 and Afadin can be found within the same protein complex, we immunoprecipitated endogenous Afadin in Caco-2 cells, a colorectal cancer cell line with strong epithelial characteristics that retains the ability to polarise. ASPP2 co-immunoprecipitated with Afadin, indicating that they are indeed found in the same protein complex (Figure 6D). To further investigate where this complex might form, we analysed the localisation of endogenous ASPP2 and Afadin in Caco-2 and MDCK cells using super-resolution Airyscan microscopy. We found the proteins colocalised primarily at tricellular junctions, where F-actin was also enriched, including in dividing cells in metaphase (Figure 6E and S5C). Their expression pattern also partially overlapped at bicellular junctions (Figure S5C). Interestingly, Afadin was also found at the mitotic spindles (Figure 6E) and cleavage furrow (Figure S5D), where ASPP2 was juxtaposed with Afadin. In E6.5 embryos, Afadin showed a similar localisation pattern to ASPP2, at the apical junction of cells in the epiblast and VE (Figure 6F and S5E). In the epiblast, similarly to ASPP2, it was more abundant at the F-actin-rich tricellular junctions.

Together, these results highlight the importance of the localisation pattern of ASPP2 in the epiblast, suggesting that it may be able to interact with F-actin at the apical junctions via its interaction with Afadin. They also suggest that this interaction might be important in dividing cells. To test this possibility, we generated *ASPP2*^RAKA/RAKA^ embryos in a C57BL/6 background carrying a LifeAct-GFP transgene^33^ that allowed us to visualise F-actin in living embryos with time-lapse confocal microscopy (Figure 6G and H and Figure S5F). During cell division events, following mitotic rounding, apical F-actin localisation was disrupted in *ASPP2*^RAKA/RAKA^ embryos whereas it was maintained in wild type embryos (Figure 6G). This was followed by an ectopic accumulation of cells at the apical surface of the epiblast, reducing the size of the proamniotic cavity (Figure 6H). These results suggest that the PP1 regulatory function of ASPP2 is required to maintain the architecture of apical F-actin, particularly during cell division events in the epiblast.

### ASPP2 supports tissue integrity across a variety of pseudostratified epithelia

Next, we investigated whether ASPP2 was only required in the epiblast or whether it also functioned in other tissues undergoing morphogenesis. To this end, we examined *ASPP2*^ΔE4/ΔE4^embryos at later stages of development around the time of gastrulation. At E7.5, during late primitive streak stages, mesoderm formation (marked by expression of T) and migration were broadly comparable between *ASPP2*^ΔE4/ΔE4^ embryos and wild type littermates, despite the absence of a proamniotic cavity and the dramatic accumulation of cells now filling the entirety of the space inside the embryos (Figure 7A, Movie 3). Furthermore, there was no difference in the velocity, directionality and distance travelled by mesoderm cells migrating from wild type and *ASPP2*^ΔE4/ΔE4^ mesoderm explants (Figure S6A and Movie 4). This suggested that ASPP2 was not required for mesoderm specification or migration.

**Figure 7:**
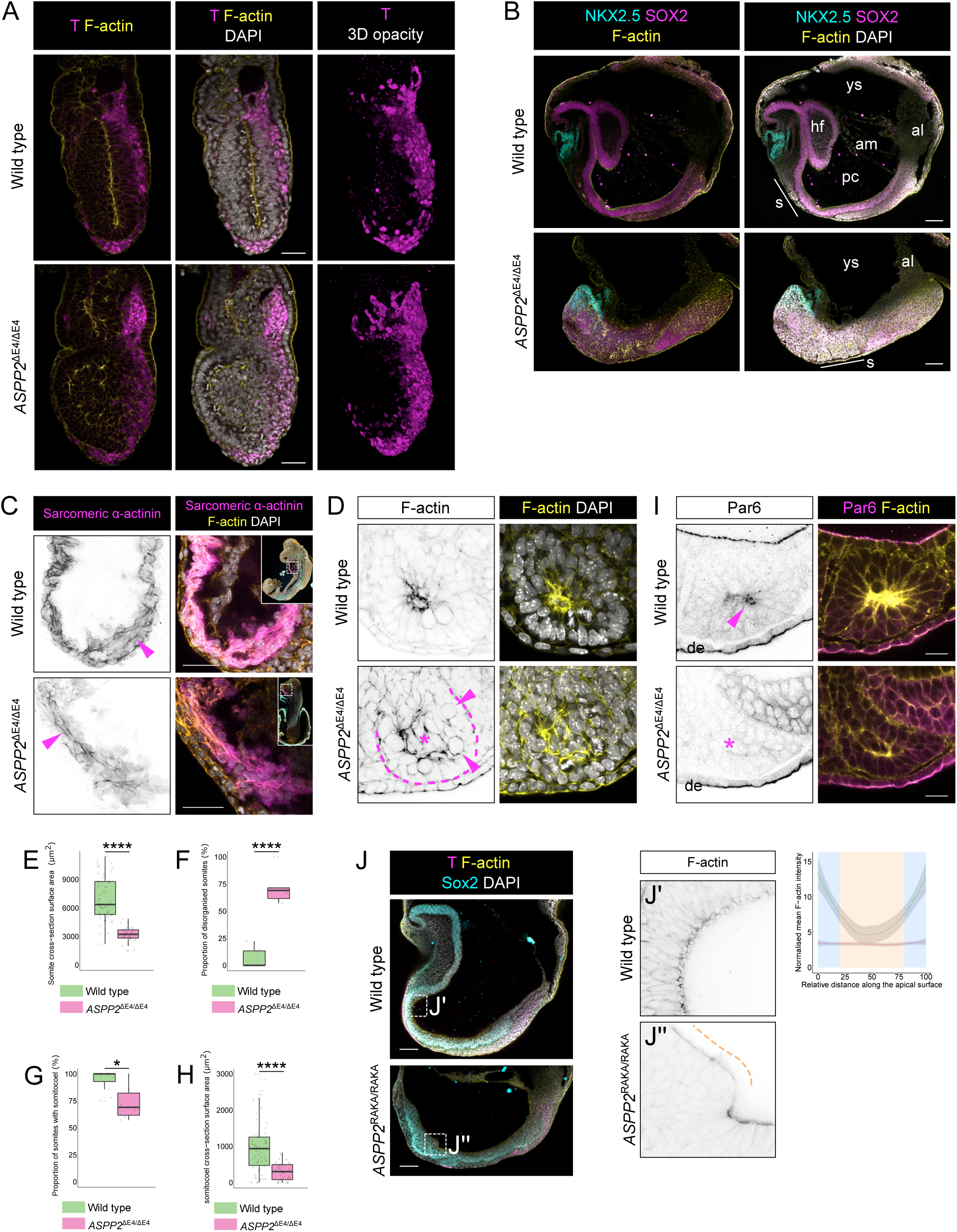
The ASPP2/PP1 complex is required for tissue integrity across a variety of highly proliferative pseudostratified epithelia. A. The primitive streak expands comparatively in E7.5 wild type and *ASPP2*^ΔE4/ΔE4^ embryos. Mesoderm cells were labelled by immunofluorescence using an antibody against Brachyury (T). B. Patterning proceeds normally in the absence of ASPP2. The ectoderm and cardiac progenitors were labelled in E8.5 wild type and *ASPP2*^ΔE4/ ΔE4^ embryos with antibodies against SOX2 and NKX2.5 respectively. ys: yolk sack, al: allantois, s: somites, hf: head fold, am: amnion, pc: proamniotic cavity. C. Cardiac progenitors can differentiate into cardiomyocytes in E9.5 *ASPP2*^ΔE4/ΔE4^ embryos. The presence of the contractile machinery (magenta arrowheads) was assessed in wild type and *ASPP2*^ΔE4/ΔE4^ embryos using an antibody against sarcomeric α-actinin. D. Somite architecture is disrupted in *ASPP2*^ΔE4/ΔE4^ embryos. The dotted line highlights the contour of a somite in an *ASPP2*^ΔE4/ΔE4^ embryo. The star indicates the ectopic accumulation of cells in the centre of this somite. Arrowheads point to mitotic figures. E-H. Quantification of somite characteristics in wild type (n=10 embryos, 58 somites) and *ASPP2*^ΔE4/ΔE4^ (n=6 embryos, 35 somites) embryos at E8.5. * p<0.05, ****p<0.0001 (Student’s T-test). I. Apical-basal polarity is defective in the somites of *ASPP2*^ΔE4/ΔE4^ embryos. Par6 localised apically in wild type somites (arrowhead) whereas it was absent in *ASPP2*^ΔE4/ΔE4^ embryos (star). de: definitive endoderm. J. Head fold formation is defective in *ASPP2*^RAKA/RAKA^ embryos. The organisation of apical F-actin was disorganised locally in the anterior ectoderm of *ASPP2*^RAKA/RAKA^ embryos (orange dotted line). F-actin signal intensity along the apical surface of ectoderm cells in disrupted areas in *ASPP2*^RAKA/RAKA^ embryos (n=3 embryos, 5 cells per embryo) was compared to wild type cells (n=3 embryos, 5 cells per embryo). Measurements were made on cross sections along the apical domain of individual ectoderm cells from apical junction to apical junction (represented with a blue background in the graph). Nuclei and the F-actin cytoskeleton were visualised with DAPI and Phalloidin respectively. Scale bars: 20 μm (D, I), 50 μm (A, C), 100 μm (B, J).

To test whether further patterning of the mesoderm occurred in the absence of ASPP2, we examined *ASPP2*^ΔE4/ΔE4^ embryos at E8.5. Morphologically, these embryos were severely disrupted, shorter along the anterior-posterior axis and without head folds (Figure 7B). However, using the cardiac progenitor marker NKX2.5, we found that this population of cells was able to migrate rostrally despite the dramatic morphological defects present in *ASPP2*^ΔE4/ΔE4^ embryos (Figure 7B). At E9.5, some cells in the anterior of these embryos were also positive for alpha sarcomeric α-actinin, suggesting that the NKX2-5 positive cells could differentiate into cardiomyocytes (Figure 7C and S6B). Next, we analysed whether the mesoderm could go on to form structurally normal somites. Using FOXC2 as a marker of somitic mesoderm, we were able to identify distinct somite-like structures in *ASPP2*^ΔE4/ΔE4^ embryos (Figure 7D and S6B). However, we found that these somites were smaller than normal (Figure 7E). Compared to wild-type control somites, a large proportion of mutant somites exhibited disrupted epithelial organisation (Figure 7D and 7F) and a reduced proportion formed cavities (Figure 6D and 6G). The relative size of the somitocoel in *ASPP2*^ΔE4/ΔE4^ embryos was also significantly reduced in comparison to wild type controls (Figure 7H). Importantly, somites with disrupted epithelial organisation and lacking cavities had features reminiscent of *ASPP2*^ΔE4/ΔE4^epiblasts: cells could be seen accumulating in the centre resulting in the obliteration of the somitocoel (Figure 7D). These cells also displayed a lack of apical Par6, suggesting that as in the epiblast, in the forming somites, apical-basal polarity was disrupted (Figure 7I).

These results suggest that ASPP2 is required not only in the epiblast, but more generally in pseudostratified epithelia^14^. To investigate this further, we tested the requirement for ASPP2 in lumen formation during cystogenesis. We derived embryonic stem cells (ESC) from embryos with exon 4 of *ASPP2* flanked by two LoxP sites. To generate *ASPP2*^ΔE4/ΔE4^ ESC, they were infected with a CRE-recombinase-expressing adenovirus. When grown in Matrigel, we found that the majority of *ASPP2*^ΔE4/ΔE4^ ESC-derived cysts failed to form lumens in comparison to control cysts (Figure S6C). Similarly, ESC derived from *ASPP2*^RAKA/RAKA^ embryos failed to form lumens in comparison to wild type ESC, suggesting that the formation of lumens during cystogenesis requires ASPP2/PP1 interaction (Figure S6D).

Since our data suggests that ASPP2 is required in regions undergoing increased mechanical stress (Figure 5A and B), we wanted to examine further the potential importance of ASPP2 during head fold formation. We took advantage of the *ASPP2*^RAKA/RAKA^ embryos in a BALB/c background as their phenotype is milder and they develop a proamniotic cavity (Figure 4A), reducing the likelihood that phenotypes observed in the head fold region are secondary defects due to overall tissue disorganisation. At E8.5, wild type embryos exhibited fully formed head folds (Figure 7J). In contrast, *ASPP2*^RAKA/RAKA^ embryos failed to form head folds in the rostral region of the ectoderm. In this region, the epithelium buckled locally, without being able to fully complete head fold morphogenesis. Interestingly, this was accompanied by a loss of organisation of F-actin at the apical junction, similarly to what was observed in the proamniotic cavity of *ASPP2*^ΔE4/ΔE4^ embryos and in the primitive streak of *ASPP2*^RAKA/RAKA^ embryos in a BALB/C background (Figure 7J). These results strongly suggest that ASPP2 is required to maintain tissue integrity by regulating F-actin organisation at the apical junctions as tensions increase in the rostral region of the ectoderm during head fold formation. Together, these results also reinforce the idea that ASPP2, and its interaction with PP1, are required during morphogenetic events that result in increased tensions at the level of apical junctions in epithelial tissues.

## DISCUSSION

Our study unveils a central role for ASPP2 in maintaining pseudostratified epithelial integrity under increased mechanical stress during major morphogenetic events: in the formation of the proamniotic cavity, the primitive streak during gastrulation, somite formation and in the head fold region.

Strikingly, ASPP2 controls proamniotic cavity formation via a mechanism that, following cell divisions, specifically prevents the most apical daughter cells from delaminating apically. ASPP2 achieves this by maintaining the integrity and organisation of the F-actin cytoskeleton at the apical surface of dividing cells. This mechanism is consistent with ASPP2 playing a role in maintaining epithelial integrity under increased mechanical stress considering that mitotic rounding results in forces sufficient to contract the epithelium in the apical-basal axis and mechanically contribute to expansion of the lumen^2^. However, it remains unclear why it is only the more apically localised daughter cells that are affected in ASPP2 mutant embryos. One possibility is that these daughters do not inherit the basal process that tethers cells to the basement membrane in pseudostratified epithelia. However, the idea that the basal process is inherited asymmetrically is controversial^34^. Other mechanisms, such as the extent of cell-cell interactions between neighbouring cells and apical daughter cells in comparison to the more basal daughters may be involved.

It has been suggested that the purpose of interkinetic nuclear migration is to ensure that cells divide apically to safeguard the integrity of pseudostratified epithelia^35^. Here we show that this is not sufficient to maintain tissue integrity as, in the absence of ASPP2, the apical-basal movement of nuclei and the position of cell divisions during IKNM proceeds unhindered. Our observations also suggest that the organisation of the F-actin cytoskeleton at the apical junctions is not required for nuclear movement during INKM and that intact basolateral domain and attachment to the basement membrane are sufficient instead.

Given that the ASPP2 interactors Par3^36,37^ and Afadin^38–40^, tricellular junctions^41^, as well as tissue tension^42–45^ have all been shown to determine cell division orientation to some extent, it was important to explore whether ASPP2 could play a role in this process. However, our results indicate that ASPP2 does not control the orientation of cell divisions in the epiblast. Similarly to previous work^26^, we also find no bias towards cell divisions orientated in the plane of the epithelium, suggesting that, at these early stages of development, planar cell polarity may not play a role in directing cell division orientation. We therefore cannot rule out that, in a different context, ASPP2 might control cell division orientation in conjunction with Afadin. In fact, later in development in E8.5 *ASPP2*^RAKA/RAKA^embryos, cells sometimes delaminated basally in the anterior regions of the ectoderm, reminiscent of the phenotype observed in SCRIB- and DLG-depleted Drosophila wing discs, where cell divisions are normally orientated in the plane of the epithelium by cell-cell junctions to maintain epithelial integrity^46^.

Our results highlight the previously underappreciated localisation of ASPP2 at tricellular junctions in epithelial cells, in particular of the epiblast, and reveal the unknown biological function of ASPP2/PP1 complexes in the regulation of F-actin organisation at the apical junction. Considering that Afadin regulates the architecture of tricellular junctions in response to tensions^31,47^, the interaction between Afadin and ASPP2 strongly suggests that ASPP2 may exert its F-actin function at tricellular junctions via Afadin. The role of Afadin in regulating the linkage between F-actin and junctions during apical constrictions48 suggests that ASPP2 may also be important in this process, which may be particularly relevant in the primitive streak. Tricellular junction are emerging as a particularly important aspect of tissue homeostasis, at the intersection between actomyosin contractility and apical-basal organisation in the context of tissue tensions^49^. We therefore suggest that ASPP2 may be directly involved in the response to tissue tension by interacting with Afadin at the level of tricellular junctions to maintain F-actin organisation.

Our study also suggests that the interaction between ASPP2 and PP1 might be essential to the well documented tumour suppressor function of ASPP2^50–52^. Simply abrogating the ability of ASPP2 to recruit PP1 is enough to induce the formation of abnormal discrete clusters of cells in the epiblast reminiscent of tumours. This suggests that mutations in ASPP2 that interfere with its interaction with PP1 might, in conjunction with mechanical stress, lead to tumour development. These mutations could be in the canonical PP1-binding domain of ASPP2, but also in other key domains which have been shown to contribute to the interaction^53^. Recent findings support the idea that ASPP2 mutations could lead to tumorigenesis in the presence of mechanical stress. Using insertional mutagenesis in mice with mammary-specific inactivation of Cdh1, ASPP2 was identified as part of a mutually exclusive group containing three other potential tumour suppressor genes (Myh9, Ppp1r12a and Ppp1r12b), suggesting that these genes target the same process^54^. With our finding that ASPP2 controls the organisation of the F-actin cytoskeleton, it now becomes apparent that, in addition to three of these genes being PP1-regulatory subunits, all four are in fact F-actin regulators. Biological studies to test specific mutations found in ASPP2 in cancer and elucidating the substrates and specific phospho-residues targeted by the ASPP2/ PP1 complex will therefore provide new insights into the tumour suppressor role of ASPP2 and might help developing new approaches to cancer treatment.

## MATERIALS AND METHODS

### Mouse strains and embryo generation

All animal experiments complied with the UK Animals (Scientific Procedures) Act 1986, were approved by the local Biological Services Ethical Review Process and were performed under UK Home Office project licenses PPL 30/3420 and PCB8EF1B4.

All mice were maintained on a 12-hour light, 12-hour dark cycle. Noon on the day of finding a vaginal plug was designated 0.5 dpc. For preimplantation stages, embryos were flushed using M2 medium (Sigma M7167) at the indicated stages. For post-implantation stages, embryos of the appropriate stage were dissected in M2 medium with fine forceps and tungsten needles.

We originally obtained *ASPP2* mutant mice in which exons 10–17 were replaced with a neo-r gene^23^from Jackson Laboratory. After careful characterisation of this mouse line, we found that the Neo cassette was not inserted in the ASPP2 locus. As a consequence, we used a different strategy to generate *ASPP2* mutant mice. C57BL/6N-Trp53bp2<tmIa (EUCOMM) heterozygous sperm (obtained from the Mary Lyon Centre) was initially used to fertilise ACTB:FLPe B6J homozygous oocytes (Jackson Laboratory). This resulted in the removal by the flippase of the LacZ and neo-r region flanked by FRT sites and the generation of heterozygous mice with one allele of ASPP2 in which exon 4 was flanked by LoxP sites. Those mice were bred in a C57BL/6J background for over four generations to breed out the rd8 mutation in the CRB1 gene found in the C57BL/6N background and eliminate the remaining FRT site left behind. They were then crossed to generate mice homozygous for the ASPP2 conditional allele in a C57BL/6J background (*ASPP2*^flE4/flE4^ mice). These mice were also crossed with Sox2Cre mice^55^ to generate mice with Exon 4 excised in one allele of *ASPP2* (*ASPP2*^WT/ ΔE4^ mice). *ASPP2*^WT/ΔE4^ mice were subsequently backcrossed into wildtype C57BL/6J mice to segregate out the Sox2Cre transgene.

*ASPP2*^WT/ΔE4^ mice were used to generate *ASPP2*^ΔE4/ΔE4^ embryos. To produce epiblast-specific *ASPP2*-null embryos (*ASPP2*^EpiΔE4/ΔE4^ embryos), *ASPP2*^WT/ΔE4^ mice homozygous for the Sox2Cre transgene were crossed with *ASPP2*^flE4/flE4^ mice. To generate *ASPP2*^ΔE4/ΔE4^ embryos with fluorescently labelled membranes, we established *ASPP2*^WT/ΔE4^mice heterozygous for the mT/mG transgene^56^ and crossed them with *ASPP2*^WT/ΔE4^ mice.

The *ASPP2*^WT/RAKA^ mice were made by inGenious Targeting Labs (Ronkonkoma, NY). A BAC clone containing exon 14 of the trp53bp2 gene was subcloned into a ∼2.4kb backbone vector (pSP72, Promega) containing an ampicillin selection cassette for retransformation of the construct prior to electroporation. A pGK-gb2 FRT Neo cassette was inserted into the gene. In the targeting vector, the wild type GTG AAA TTC was mutated to GCG AAA GCC by overlap extension PCR and introduced into C57BL/6 × 129/SvEv ES cells by electroporation. Inclusion of the mutations in positive ES cell clones was confirmed by PCR, sequencing and Southern blotting. ES cells were microinjected into C57BL/6 blastocysts and resulting chimeras mated with C57BL/6 FLP mice to remove the Neo cassette. The presence of the mutation was confirmed by sequencing. Mice were then back-crossed with BALB/cOlaHsd or C57BL/6J mice for at least eight generations to obtain the RAKA mutation in the respective pure background. *ASPP2*^RAKA/RAKA^ embryos were generated from heterozygous crosses. To generate LifeAct-GFP-positive *ASPP2*^RAKA/RAKA^ embryos, we generated *ASPP2*^WT/ RAKA^ mice heterozygous for the LifeAct-GFP transgene^33^.

### siRNA microinjections

siGENOME RISC-Free Control siRNA (Dharmacon) and Silencer Select Pre-designed siRNAs against mouse ASPP2 (Ambion) were resuspended in nuclease-free sterile water and used at 20 μM. For zygotes, 3 to 4 week old CD-1 females (Charles River UK) were injected intraperitoneally with 5 IU of PMSG (Intervet) and 48 h later with 5 IU of hCG (Intervet), and were paired with C57Bl/6J male mice (in house). Zygotes were retrieved from oviductal ampullae at 20 hours post-hCG. Cumulus-enclosed zygotes were denuded by exposure to 1 mg/mL hyaluronidase (Sigma) in modified mHTF (Life Global) containing 3mg/ml BSA for 3–6 min and cultured in LGGG-020 (life Global) containing 3mg/ml BSA in the presence of 5% CO2 at 37°C. Microinjection of zygotes commenced 2 hours after release from cumulus mass. Zygotes with a normal morphology were microinjected into the cytoplasm in 30µl drops of modified HTF media containing 4mg/ml BSA using a PMM-150FU Piezo impact drive (Primetech) using homemade glass capillaries with 5–10 pL of siRNA. Zygotes were returned to LGGG-020 containing 3mg/ml BSA in the presence of 5% CO2 at 37°C until analysis.

### Human embryo collection

Human embryos were donated from patients attending the Oxford Fertility with approval from the Human Fertilization and Embryology Authority (centre 0035, project RO198) and the Oxfordshire Research Ethics Committee (Reference number 14/SC/0011). Informed consent was attained from all patients. Embryos were fixed in 4% paraformaldehyde, washed twice and kept in PBS containing 2% bovine serum albumin (PBS-BSA) at 4°C until they were used for immunohistochemistry.

### Wholemount immunohistochemistry

Post-implantation embryos were fixed in 4% paraformaldehyde in phosphate-buffered saline (PBS) at room temperature for 20 to 45 minutes depending on embryo stages. Embryos were washed twice for 10 minutes in 0.1% PBS-Tween (PBS containing 0.1% Tween 20). Embryos were then permeabilized with 0.25% PBS-Triton (PBS containing 0.25 Triton X-100) for 25 minutes to 1 hour depending on embryo stages and then washed twice for 10 minutes in 0.1% PBS-Tween. Embryos were incubated overnight in a blocking solution (3% Bovine serum albumin, 2.5% donkey serum in 0.1% PBS-Tween). The next day, primary antibodies were diluted in blocking solution and added to the embryos overnight. The following day, embryos were washed three times for 15 minutes in 0.1% PBS-Tween and then incubated with secondary antibodies and Phalloidin diluted in blocking solution overnight. Finally, embryos were washed four times in 0.1% PBS-Tween and kept in DAPI-containing VECTASHIELD Antifade Mounting Medium (Vector Laboratories) at 4°C until used for imaging. Short incubation steps were carried out in wells of a 12-well plate on a rocker at room temperature and overnight steps were carried out in 1.5 mL Eppendorf tubes at 4°C.

For preimplantation embryos, fixation and permeabilization times were reduced to 15 minutes and 2% PBS-BSA (PBS containing 2% Bovine serum albumin) was used for washing steps. Blocking and secondary antibody incubation steps were reduced to one hour. Embryos were transferred between solutions by mouth-pipetting. the embryos were mounted in 8-well chambers in droplets consisting of 0.5 μL DAPI-containing VECTASHIELD and 0.5 μL 2% PBS-BSA. After mounting the embryos were kept in the dark at 4°C until they were imaged.

### Immunocytochemistry

Caco-2 and MDCK.2 cells were maintained in Dulbecco’s modified Eagle’s medium containing 10% fetal bovine serum, penicillin, and streptomycin at 37°C in a 5% CO2 atmosphere incubator. In preparation for immunocytochemistry, Caco-2 cells were seeded onto coverslips in 24-well plates with fresh medium. 48 hours later, cells were fixed with 4% paraformaldehyde (in PBS) for 10 minutes, washed twice in PBS, and then permeabilized with 0.1% Triton X-100 in PBS for 4 minutes. Cells were washed twice in PBS and 2% PBS-BSA was then used as a blocking solution for 30 minutes prior to incubation with primary antibodies. Primary antibodies were diluted in 2% PBS-BSA and applied to cells for 40 minutes. Cells were then washed three times with PBS. Secondary antibodies (1:400), DAPI (1:2000, Invitrogen) and Phalloidin (1:400) were diluted in 2% PBS-BSA and applied to cells for 20 minutes. Coverslips were then washed three times with PBS and mounted onto glass slides with a small drop of Fluoromout-G (SouthernBiotech). They were air-dried before being sealed with nail varnish. All incubation steps were carried out at room temperature on a rocker. Samples were kept in the dark at 4°C until they were imaged.

### Antibodies and phalloidin conjugates

The following antibodies were used at the stated dilutions: rabbit anti-ASPP2 (Sigma, HPA021603), 1:100-1:200 (IHC); mouse anti-ASPP2 (Santa Cruz Biotechnologies, sc135818), 1:100 (ICC), 1:1000 (IB); mouse anti-YAP (Santa Cruz Biotechnology, sc-101199), 1:100 (IHC); rabbit anti-pYAP S127 (Cell Signaling, 4911), 1:100 (IHC); rabbit anti-Par3 (Millipore, 07-330), 1:100 (IHC); rabbit anti-Pard6b (Santa Cruz Biotechnology, sc-67393), 1:100 (IHC); rabbit anti-SCRIB (Santa Cruz Biotechnology, sc28737), 1:100 (IHC); goat anti-Brachyury (Santa Cruz Biotechnology, sc17745), 1:100 (IHC); rabbit anti-Sarcomeric α-actinin (Abcam, ab68167), 1:100 (IHC); mouse anti-FOXC2 (Santa Cruz Biotechnology, sc515234), 1:100 (IHC); rabbit anti-SOX-2 (Millipore, AB5603), 2 μL per mg of cell lysate (co-IP), 1:100 (IHC); goat anti-NKX2.5 (Santa Cruz Biotechnology, sc8697), 1:100 (IHC); rabbit anti-Afadin (Sigma, A0224), 2 μL per mg of cell lysate (co-IP), 1:100 (IHC, ICC), 1:1000 (IB); rabbit anti-Laminin (Sigma, L9393), 1:200 (IHC); goat anti-AMOT (Santa Cruz Biotechnologies, sc82491), 1:200 (IHC); goat anti-GATA-6 (R&D Systems, AF1700), 1:100 (IHC); rabbit anti-Myosin IIa (Cell Signaling, #3403), 1:100; rabbit anti-phospho-Myosin light chain 2 (Cell Signaling, #3674), 1:100. The following were used at 1:100 for IHC and 1:400 for ICC: Alexa fluor 555 donkey-anti-mouse (Invitrogen, A-31570), Alexa fluor 647 goat-anti-rat (Invitrogen, A-21247), Alexa fluor 488 donkey-anti-rabbit (Invitrogen, A21206), Phalloidin-Atto 488 (Sigma, 49409), Phalloidin–Atto 647N (Sigma, 65906).

### Confocal microscopy, image analysis and quantification

Samples were imaged on a Zeiss Airyscan LSM 880 confocal microscope with a C-Apochromat 40x/1.2 W Korr M27 water immersion objective or a Plan-Apochromat 63x/1.4 OIL DIC M27 objective. For super-resolution imaging, an Airyscan detector was used^57^. Volocity (version 6.3.1, PerkinElmer) and Zen (Zeiss) software were used to produce maximum intensity projections and 3D opacity renderings. Image analysis was performed on optical sections. For signal intensity profiles along the apical-basal axis and across tricellular junctions, the arrow tool in the Zen software was used. Anterior and posterior embryo widths measurements were made using the line tool in Volocity.

For F-actin signal intensity profiles across the apical surface of epiblast or ectoderm cells, Fiji’s freehand line tool with a width of “3” was used. Because the size of the apical domain was different for each cell measured, distances were expressed as percentages, with 100% representing the total distance across the apical domain. To account for depth-dependent signal attenuation, F-actin signal intensity at the apical domain was normalized by mean F-actin intensity in the nucleus of the cell measured. In each experiment, for each genotype, three embryos were used for measurements and 5 cells were analysed per embryo. The LOWES method was used to fit a line to the data.

### Mouse embryo culture for live imaging and image analysis

To restrain embryo movement during imaging, lanes were constructed inside 8-well Lab-Tek II chamber slide (Nunc), using glass rods made from hand-drawn glass capillaries. Shorter pieces were used as spaces between two rods to create a space slightly wider than an embryo. Silicone grease was used to maintain the rods together. Each well was filled with medium containing 50% phenol red-free CMRL (PAN-Biotech, Germany) supplemented with 10 mM L/glutamine (Sigma-Aldrich) and 50% Knockout Serum Replacement (Life Technologies, England). The chamber was equilibrated at 37°C and an atmosphere of 5% CO2 for at least 2 h prior to use. Freshly dissected embryos were placed in the lanes between two rods and allowed to settle prior to imaging on a Zeiss LSM 880 confocal microscope equipped with an environmental chamber to maintain conditions of 37°C and 5% CO2. Embryos were imaged with a C-Apochromat 40x/1.2 W Korr M27 water immersion objective. Using a laser excitation wavelength of 561 nm, embryos labelled with mT/mG were imaged every 7.5 minutes and for each time point, nine z-sections were acquired every 3 μm around the midsagittal plane for up to 10 hours. For LifeAct-positive embryos, a laser excitation wavelength of 488 nm was used, and embryos were imaged every 15 minutes for 6 hours. For each time point, 12 z-sections every 1.5 μm were collected around the midsagittal plane.

Daughter cell movement was quantified using the Fiji plugin TrackMate (v5.2.0). Timepoints were registered using Fiji. Jittering was accounted for by correcting cell coordinates relatively to the centre of the embryonic region. The distance travelled by daughter cells (d) was analysed by calculating the distance between the coordinates of their final position and the coordinates of their respective mother cell immediately prior cell division. The direction of daughter cell movement (θ) was analysed by calculating the angle between the vector describing cell movement (that is the vector originating from the coordinates of the mother cell immediately prior cell division to the coordinates of the daughter cell at its final position) and the vector from the coordinates of the mother cell prior cell division to the coordinates of the embryonic region’s centre. To establish the angle of cell division, we first defined a vector starting at the coordinates of one daughter and ending at the coordinates of the other immediately after cell division. We then defined a second vector originating halfway between the two daughters and terminating at the centre of the embryonic region. The angle of cell division was defined as the angle between those two vectors. The relative position of cell divisions was defined as the distance between the position of the mother cell immediately prior to cell division and the base of the epiblast.

### Embryo culture in channels

Channels were formed by casting a 5% (which corresponded to approximately 4.2kPa stiffness^58^) acrylamide hydrogel (containing 39:1 bisacrylamide) around 60 µm wires within the confinement of a two-part mould (10×10×1mm). Ammonium persulphate (0.1%) and TEMED (1%) were added to polymerize polyacrylamide. The wires were then removed to form cylindrical cavities within hydrogel pieces. The hydrogels were carefully washed and equilibrated in embryo culture media at 37°C and 5% CO2. The embryos were then inserted into the channels using a glass capillary with a diameter slightly larger than the embryo itself. It was used to stretch the hydrogel channel before injecting the embryos and letting the channels relax and deform the embryos. Cell viability in channels had previously been assessed without any noticeable difference with control embryos^27^. After 30 minutes, embryos were fixed inside the hydrogels with 4% PFA for 35 minutes. Once fixed, embryos were removed from the hydrogel channels and wholemount immunohistochemistry was performed.

### Co-immunoprecipitation and SDS-PAGE/Immunoblotting

For immunoprecipitation experiments, Caco-2 cells from confluent 10 cm diameter dishes were washed twice with PBS and then lysed in 500 μL of a buffer containing 50 mM Tris-HCl at pH 8, 150 mM NaCl, 1 mM EDTA, Complete Protease Inhibitor Cocktail (Roche) and 1% Triton X-100. Lysates were left on ice for 30 minutes, briefly sonicated and spun down at 21,000 × g for 30 minutes at 4°C. The supernatant was transferred to another tube and protein concentration was measured (Bradford, Bio-Rad). 1 mg of protein lysate was used per condition. Lysates were precleared using 20 μL protein G Sepharose 4 fast flow (1:1 in PBS, GE Healthcare) for 30 minutes at 4°C on a shaker. The supernatant was incubated for 30 minutes at 4°C on a shaker with 2 μL of the indicated antibody. 30 μL protein G Sepharose 4 Fast Flow (1:1 in PBS) was added to each condition and samples were incubated overnight at 4°C on a shaker. Samples were washed 5 times with ice cold lysis buffer. 25 μL sample buffer was added and samples were incubated at 95°C for 5 minutes before being subjected to SDS-PAGE/Immunoblotting.

### Mesoderm explants and mesoderm cell migration

*ASPP2*^WT/RAKA^ mice heterozygous for the LifeAct-GFP transgene were crossed and E7.5 embryos were dissected in M2. Embryos were then incubated in a 2.5% pancreatin mixture on ice for 20 minutes. Using tungsten needles, the visceral endoderm layer was removed and then the mesodermal wings were separated from the underlying epiblast. Mesodermal tissue was grown in fibronectin-coated 8-well Lab-Tek II chamber slides and cultured in DMEM containing 10% fetal bovine serum, penicillin, and streptomycin at 37°C and 5% CO2^59^. Samples were imaged on a Zeiss LSM 880 confocal microscope equipped with an environmental chamber to maintain conditions of 37°C and 5% CO2. A laser excitation wavelength of 488 nm was used, and explants were imaged every 5 minutes for 5 hours. For each time point, 9 z-section with 1 μm step were collected.

Individual cells migrating away from the explants were tracked using the manual tracking plugin in Fiji. The movement, velocity and directionality of individual cells was analysed. Movement represented the total distance travelled in μm by an individual cell. Velocity represented the average speed in μm/min of a given cell. Directionality was used as a measure of how direct or convoluted a cell’s path was and was calculated as the ratio between the total distance travelled and the distance in a straight line between a cell’s start and end position^60^.

### Embryonic stem cell-derived cysts

Using small-molecule inhibitors of Erk and Gsk3 signalling^61^, *ASPP2*^flE4/flE4^ and *ASPP2*^RAKA/RAKA^(and *ASPP2*^WT/WT^ controls) ESC were generated from flushed E2.5 embryos obtained from crosses between *ASPP2*^flE4/flE4^ and *ASPP2*^WT/RAKA^ mice, respectively. Briefly, embryos were grown for two days in organ culture dishes, containing pre-equilibrated preimplantation embryo culture media supplemented with 1 µM PDO325901 and 3 µM CHIR99021 (Sigma-Aldrich). Embryos were grown one more day in NDiff 227 media (Takara) supplemented with 1 µM PDO325901 and 3 µM CHIR99021 (NDiff + 2i). The trophectoderm was removed by immunosurgery and “epiblasts” were grown in gelatinised dishes in the presence of NDiff + 2i and ESGRO (recombinant mouse LIF Protein, Millipore) to establish ESC lines.

*ASPP2*^flE4/flE4^ ESC were infected with an Ad-CMV-iCre adenovirus (Vector Biolabs) to delete exon 4 of *ASPP2*. Deletion of exon 4 was assessed by PCR. Non-infected *ASPP2*^flE4/flE4^ ESC were used as controls. Wild type ESC derived from litter mates were used as controls for *ASPP2*^RAKA/ RAKA^ ESC. To form cysts, 4500 ESC were resuspended in 150 μL Matrigel (354230, Corning) and plated into a well of an 8-well Lab-Tek II chamber slide. The gel was left to set for 10 minutes at 37°C before 300 μL differentiation medium (DMEM supplemented with 15% FCS, 1% Penicillin/Streptomycin, 1% Glutamine, 1% MEM non-essential amino acids, 0.1 mM 2-mercaptoethanol and 1mM sodium pyruvate) was added. ESC were grown for 72 hours at 37°C and 5% CO2 before immunostaining was performed.

## ACKNOWLEDGEMENTS

This work was funded by Wellcome Senior Investigator Award 103788/Z/14/Z (SS). We thank Jenny Nichols for advice and protocols for deriving Embryonic Stem Cells.

## AUTHOR CONTRIBUTIONS

C.R., and S.S. led the project, conceived, and designed the experiments. C.R., E.Sa., E.Sl., J.Go., N. V., K.L., J.Ga., and T.N. conducted the experiments. C.R. and S.S. analysed the data. C.R. performed the statistical analyses. E.Sl. and X.L. designed and established the *ASPP2*^WT/RAKA^ mouse line. H.H. and F.Z. performed the unwrapping of ASPP2 and F-actin immunostaining in E5.5 embryos. A.V., C.J., T.C., K.C. and C.G. organised the collection of human embryos. C.R. and S.S. wrote the manuscript.

---

**Figure S1:**
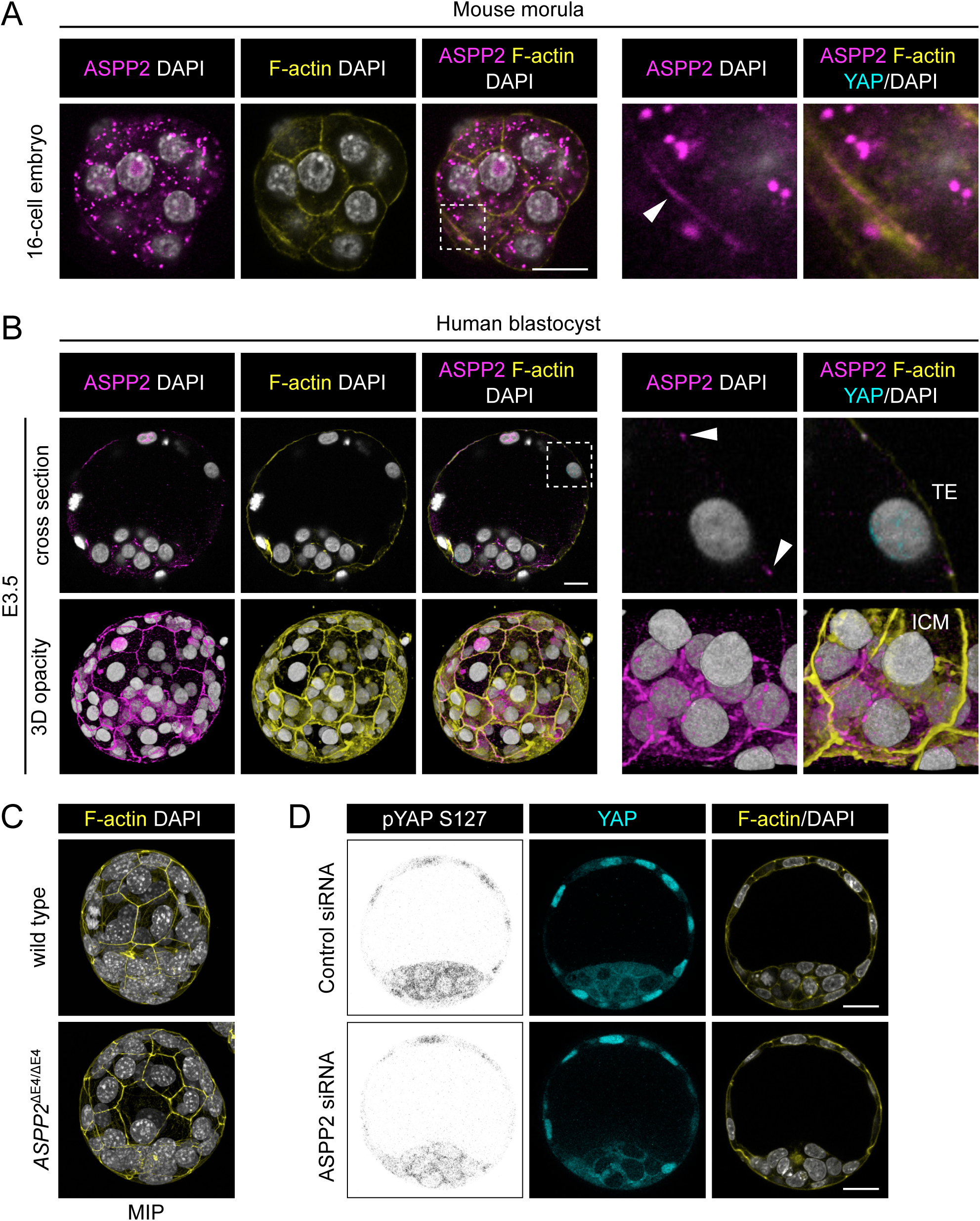
ASPP2 is not required during preimplantation development. A. Localisation pattern of ASPP2 in 16-cell embryos. A cross section through the equatorial plane of a representative embryo is shown. The F-actin cytoskeleton and nuclei were visualised using Phalloidin and DAPI respectively. The dashed area is magnified on the right. The white arrowhead shows ASPP2 and F-actin colocalising at an apical junction between two outside cells. Scale bars: 20 μm (left panel) and 5 μm (right panel). B. localisation pattern of ASPP2 in human blastocysts. The top panel shows a cross section through the equatorial plane of a representative embryo. The dashed area is magnified on the right to highlight the colocalisation between ASPP2 and F-actin at the level of apical junctions in the trophectoderm (TE) (white arrowheads). The bottom row shows a 3D opacity rendering of the same embryo in its totality (left panel) and a focus on its inner cell mass (right panel). TE: trophectoderm; ICM: Inner cell mass. Scale bar: 20 μm. C. F-actin is normally distributed at the level of apical junctions in the TE of *ASPP2*^ΔE4/ΔE4^ embryos. Maximum intensity projections of representative wild type and *ASPP2*^ΔE4/ΔE4^ embryos stained with Phalloidin and DAPI. D. ASPP2 knockdown in E3.5 embryos using siRNA targeting ASPP2 mRNA. Note that the localisation pattern of pYAP S127 was similar in control and ASPP2-depleted embryos. Scale bar: 20 μm.

**Figure S2:**
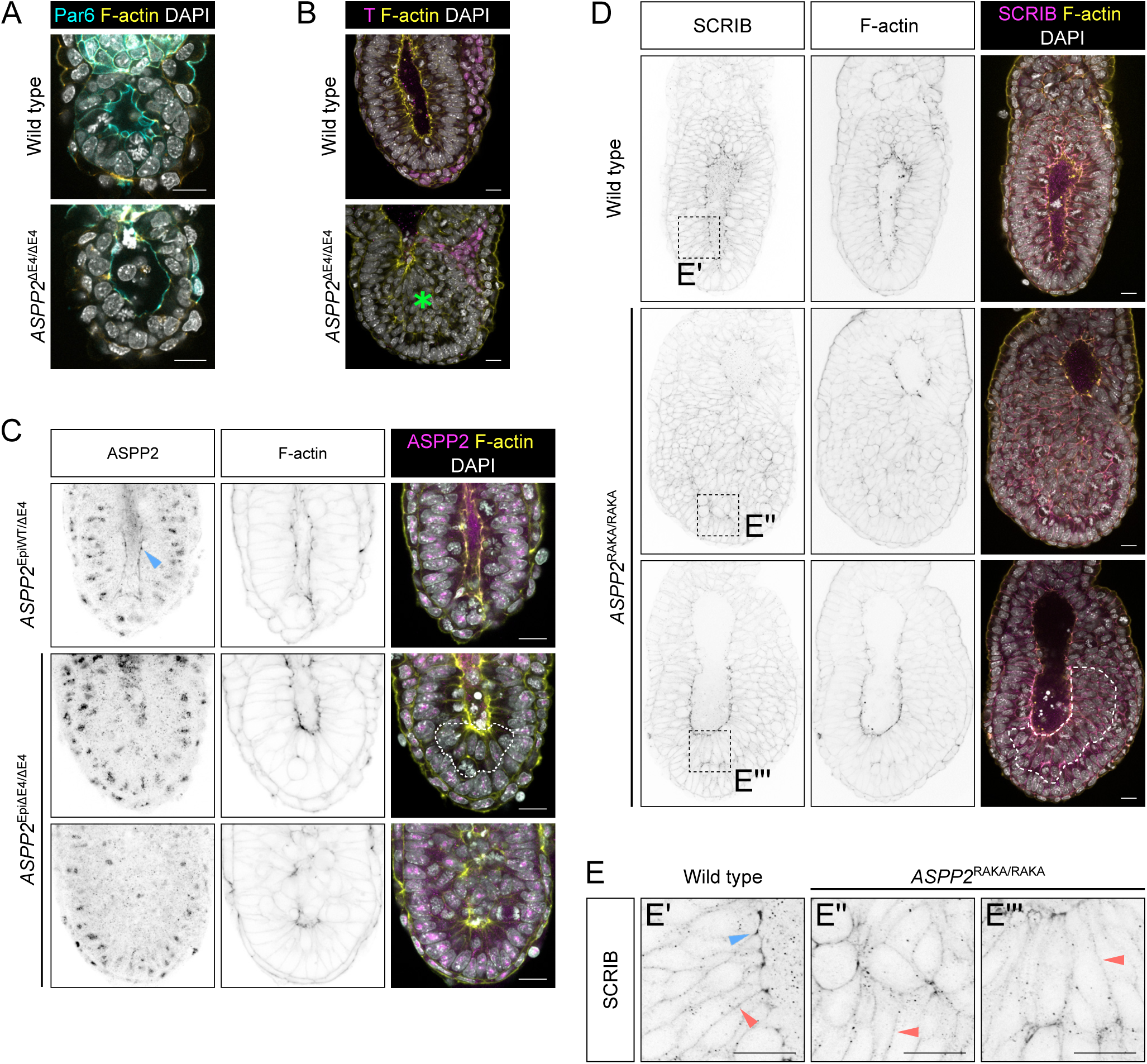
ASPP2 is required specifically in the epiblast during proamniotic cavity formation. A. Immunofluorescence of wild type and *ASPP2*^ΔE4/ΔE4^ E5.5 embryos using an anti-Par6 antibody. B. Phenotype of *ASPP2*^ΔE4/ΔE4^ embryos at E7.5. Brachyury (T) localisation was analysed by indirect immunofluorescence. The ectopic accumulation of cells in the proamniotic cavity of *ASPP2*^ΔE4/ΔE4^ embryos is indicated by a green star. C. ASPP2 expression was conditionally prevented in the epiblast to test whether ASPP2 was specifically required in the epiblast (*ASPP2*^EpiΔE4/ΔE4^ embryos). ASPP2 expression pattern was analysed by indirect immunofluorescence. ASPP2 proteins were completely absent at the apical junction of epiblast cells in *ASPP2*^EpiΔE4/ΔE4^ embryos. Note that the ASPP2 antibody results in non-specific nuclear signal (also seen in Figure 1C when depleting ASPP2 by siRNA). The dashed area highlights the ectopic accumulation of cells in the epiblast. D. SCRIB expression pattern was analysed by indirect immunofluorescence in wild type and *ASPP2*^RAKA/RAKA^ embryos. The ectopic accumulations of cells in the epiblast of *ASPP2*^RAKA/RAKA^ embryos was highlighted by a dashed line. E. Magnification of the corresponding regions in D. The blue arrowhead points to the enrichment of SCRIB at the apical junctions. Red arrowheads point to basolateral SCRIB. The F-actin cytoskeleton and nuclei were visualised using Phalloidin and DAPI respectively. Scale bars: 20 μm.

**Figure S3:**
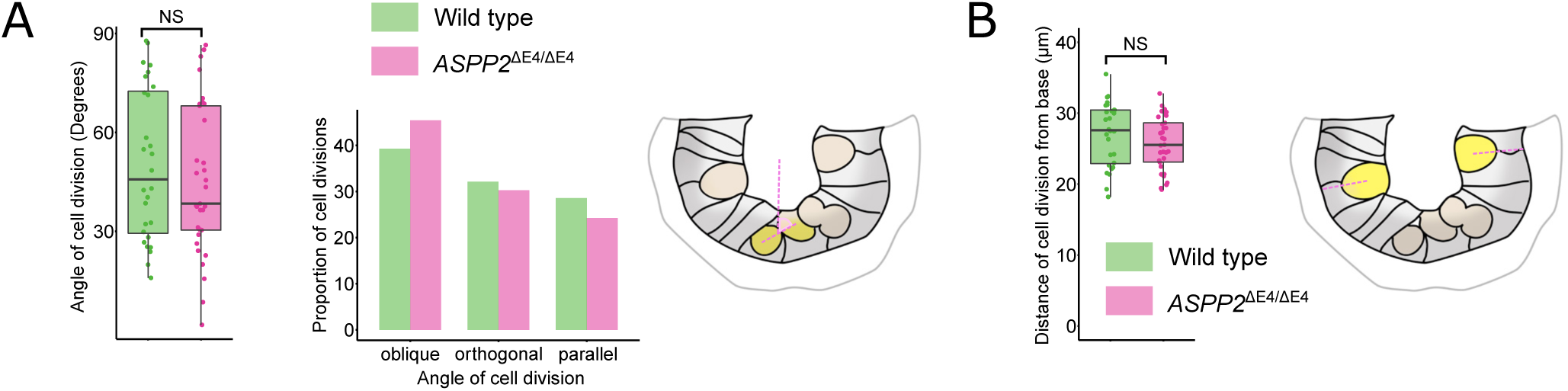
The angle of cell division and their relative position from the base of the epiblast is unaffected in the absence of ASPP2. A. Quantification of cell division angles in the epiblasts of wild type (n=3 embryos, 28 cells) and *ASPP2*^ΔE4/ ΔE4^ embryos (n=3 embryos, 33 cells). Left panel: Comparison of all cell division angles in wild type and *ASPP2*^ΔE4/ΔE4^ embryos. NS: non-significant (nested ANOVA). right panel: Cell division angles were defined as either parallel (0° to 30°), oblique (30° to 60°) or orthogonal (60° to 90°). NS: non-significant (Fisher’s exact test of independence). B. Position of cell division events in wild type and *ASPP2*^ΔE4/ΔE4^ embryos. The relative position of cell division events was expressed as the distance between mother cell position immediately prior to a division event and the base of the epithelium. NS: non-significant (nested ANOVA).

**Figure S4:**
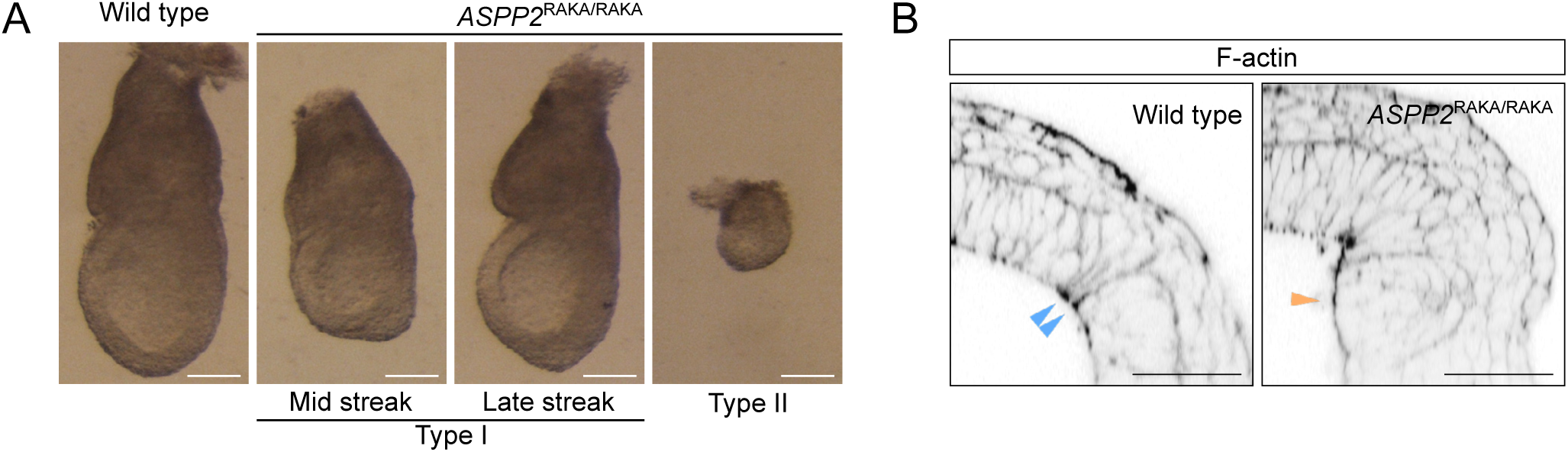
Gross defects in ASPP2^RAKA/RAKA^ embryos in a BALB/c background at E7.5. A. Bright field images of wild type and type I (34/41, 82.9%) and type II (7/41, 17.1%) phenotypes observed in *ASPP2*^RAKA/RAKA^ embryos. Type I embryos exhibited a strong accumulation of cells in their posterior. B. Cells ectopically accumulating in the primitive streak region are unable to apically constrict and do not have enriched F-actin at the apical junctions (orange arrowhead) in comparison to wild type (blue arrow heads). The F-actin cytoskeleton was visualised using Phalloidin. Scale bars: 20 μm.

**Figure S5:**
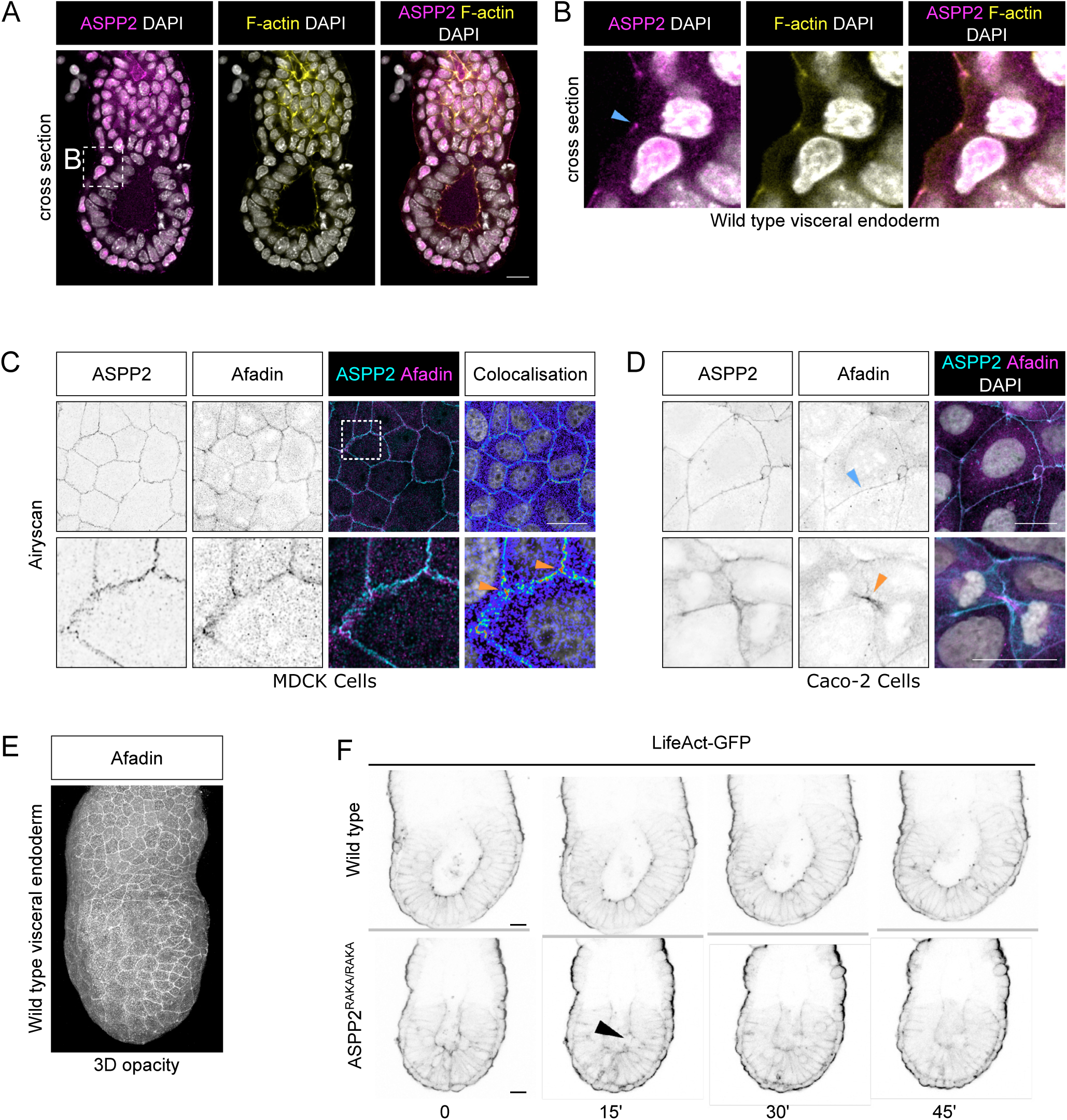
ASPP2 is required for the integrity of the F-actin cytoskeleton in the epiblast as cells divide. A. The localisation pattern of ASPP2 was analysed in wild type E5.5 embryos by immunofluorescence. B. Magnification of the boxed region in A, showing the localisation pattern of ASPP2 in the VE at the apical junctions (blue arrowhead). Note that ASPP2’s nuclear signal is non-specific. C. Super-resolution Airyscan imaging of MDCK cells immunostained for ASPP2 and Afadin. Orange arrowheads highlight the colocalisation of Afadin and ASPP2 at tricellular junctions. D. The localisation pattern of ASPP2 and Afadin was analysed in Caco-2 cells. Afadin and ASPP2, in addition to being enriched at tricellular junctions, could also be found colocalising at bicellular junctions (blue arrowhead) and were in close proximity at the cleavage furrow (orange arrowhead). E. 3D opacity rendering showing the localisation at the apical junctions of the VE in an E6.5 wild type embryo. F. Time-lapse imaging of wild type and *ASPP2*^RAKA/RAKA^ LifeAct-GFP positive embryos. Note how apical F-actin is disrupted in *ASPP2*^RAKA/RAKA^ LifeAct-GFP positive embryos following a cell division event (black arrowhead). Nuclei and the F-actin cytoskeleton were visualised with DAPI and Phalloidin respectively. Scale bars: 20 μm.

**Figure S6:**
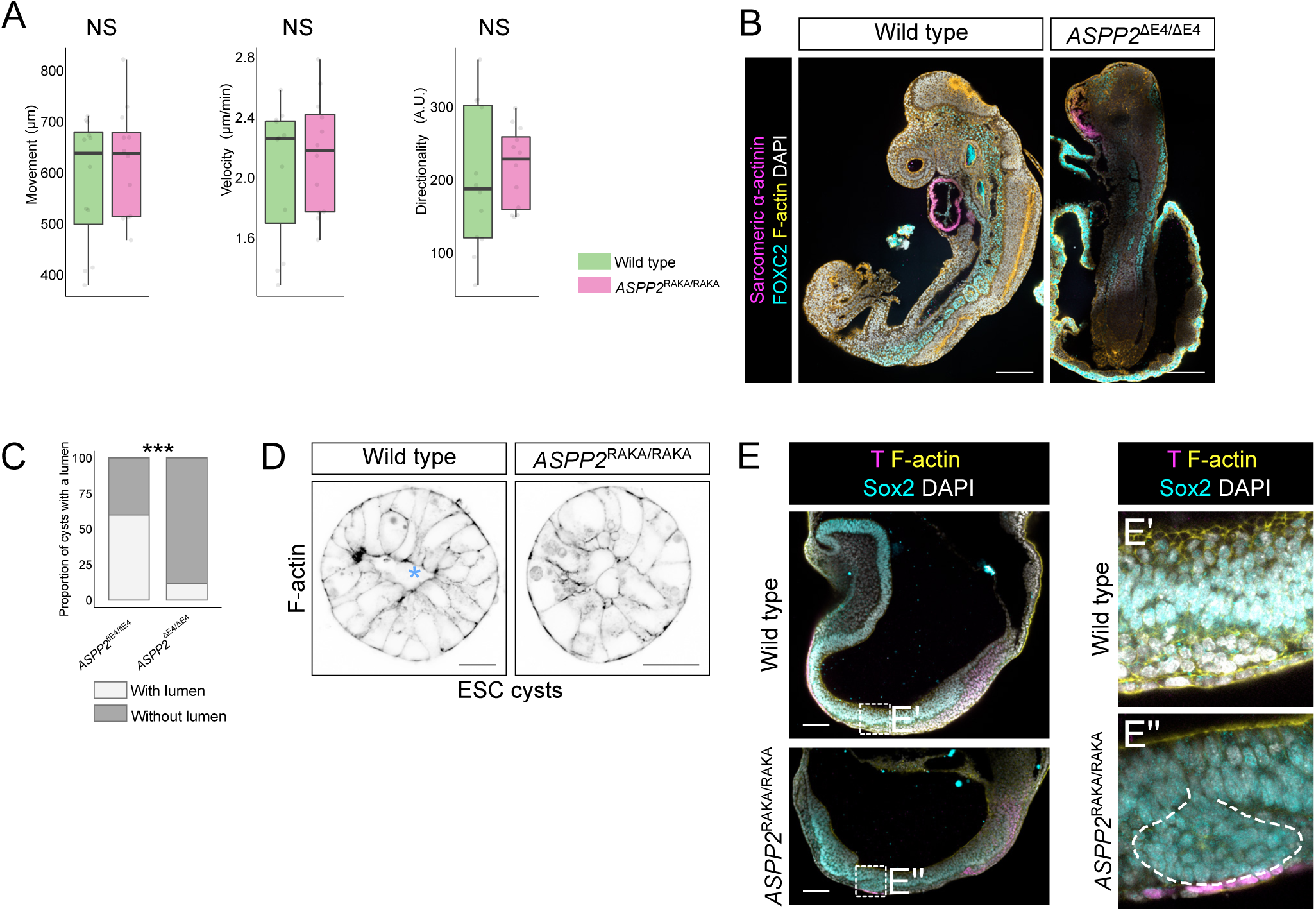
ASPP2 is required for cavity architecture in ESC cysts. A. Mesoderm cell migration is unaffected in the absence of ASPP2. Cell movement, velocity, and directionality from wild type (n=4 explants, 3 cells per explant) and *ASPP2*^RAKA/RAKA^ (n=4 explants, 3 cells per explant) mesoderm explants were quantified. NS: non-significant (nested ANOVA). B. E9.5 wild type and *ASPP2*^ΔE4/ΔE4^ embryos labelled by immunofluorescence with antibodies against FOXC2 (somitic mesoderm) and sarcomeric α-actinin (sarcomeres in cardiomyocytes). Nuclei and the F-actin cytoskeleton were visualised with DAPI and Phalloidin respectively. Scale bars: 200 μm. C. Quantification of the proportion of *ASPP2*^flE4/flE4^ and *ASPP2*^ΔE4/ΔE4^ ESC-derived cysts with lumens after three days in culture in Matrigel. D. Representative images of wild type and *ASPP2*^RAKA/RAKA^ ESC-derived cysts after three days in culture in Matrigel. The blue star indicates the presence of a cavity in the wild type cyst. E. Cells from the ectoderm occasionally delaminate basally into the underlying mesoderm in E8.5 *ASPP2*^RAKA/RAKA^ embryos. The F-actin cytoskeleton was visualised with Phalloidin respectively. Scale bars: 20 μm.

***MOVIE LEGENDS***

***Movie 1: Time lapse imaging of wild type and ASPP2***^***ΔE4/ΔE4***^ ***embryos with mT/mG-labelled cell membranes***

***Movie 2: Airyscan imaging and 3D rendering of F-actin in the primitive streak region of representative wild type and ASPP2***^***RAKA/RAKA***^ ***embryos***

***Movie 3: D rendering showing that the primitive streak expands comparatively in E7***.***5 wild type and ASPP2***^***ΔE4/ΔE4***^ ***embryos***. Mesoderm cells were labelled by immunofluorescence using an antibody against Brachyury (T). Nuclei and the F-actin cytoskeleton were visualised with DAPI and Phalloidin respectively.

***Movie 4: Time lapse imaging of wild type and ASPP2***^***RAKA/RAKA***^ ***mesoderm explants positive for the LifeAct-GFP transgene***

